# 50 years of antibody numbering schemes: a statistical and structural evaluation reveals key differences and limitations

**DOI:** 10.1101/2024.07.08.602590

**Authors:** Zirui Zhu, Katherine S. Olson, Thomas J. Magliery

## Abstract

The complementarity determining region (CDR) of antibodies represents the most diverse region both in terms of sequence and structural characteristics, playing the most critical role in antibody recognition and binding for immune responses. Over the past decades, several numbering schemes have been introduced to define CDRs based on sequence. However, the existence of diverse numbering schemes has led to potential confusion, and a comprehensive evaluation of these schemes is lacking. We employ statistical analyses to quantify the diversity of CDRs compared to the framework regions. Comparative analyses across different numbering schemes demonstrates notable variations in CDR definitions. The Kabat and AbM numbering schemes tend to incorporate more conserved residues into their CDR definitions, whereas CDRs defined by the Chothia and IMGT numbering schemes display greater diversity, sometimes missing certain loop residues. Notably, we identify a critical residue, L29, within the kappa light chain CDR1, which appears to act as a pivotal structural point within the loop. In contrast, most numbering schemes designate the topological equivalent point in the lambda light chain as L30, suggesting the need for further refinement in the current numbering schemes. These findings shed light on regional sequence and structural conservation within antibody sequence databases while also highlighting discrepancies stemming from different numbering schemes. These insights yield valuable guidelines for the precise delineation of antibody CDRs and the strategic design of antibody repertoires, with practical implications in developing innovative antibody-based therapeutics and diagnostics.

**GRAPHIC ABSTRACT:** 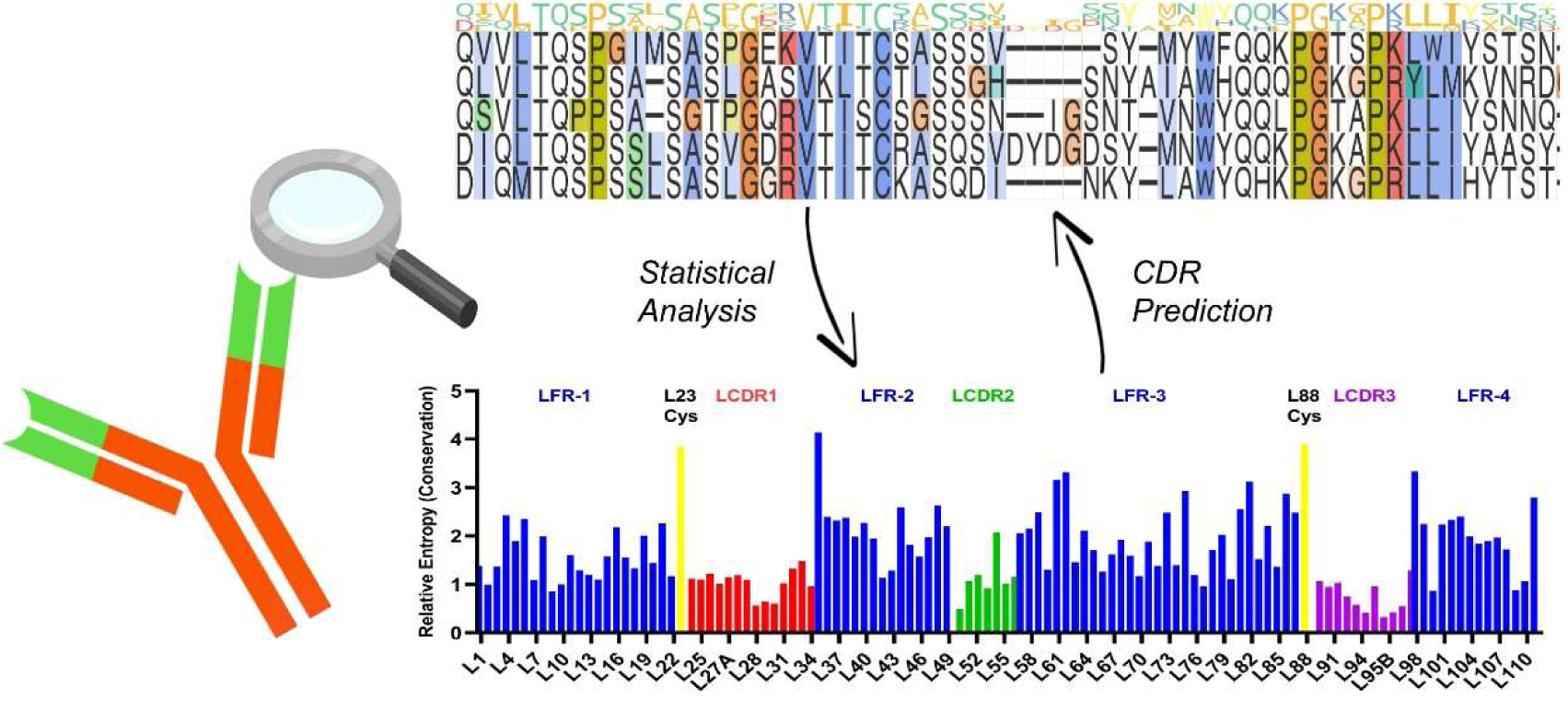

**HIGHLIGHTS:**

- Amino acid distribution analysis conducted on 124,317 sequences archived in the abYsis database showed lower conservation among CDR residues, as defined by Kabat numbering, compared to the framework region.
- Both LCDR3 and HCDR3 exhibited particularly high levels of diversity compared to the other four CDR loops, highlighting additional functional roles of the CDR3 loops relative to the others.
- Two distinct residues, L29 and H30, displayed highly conserved hydrophobic side chain characteristics and played unique conformational roles within the CDR1 loop, contributing to the formation of an ‘M-shaped’ structure. This residue divides the C-terminus CDR1 loop, which was observed to be less conserved and have greater length diversity, and the N-terminus CRD1 loop.
- The Kabat numbering scheme appears to overestimate the LCDR2 and HCDR2 loops and underestimate HCDR1, only encompassing the C-terminal loop of the ‘M-shaped’ CDR1 loop.
- Structural analysis revealed that a residue within LCDR1 was numbered differently in the two subtypes (kappa and lambda) of the variable light domain. This discrepancy suggests the need for further refinement in the current major numbering schemes.

## INTRODUCTION

Antibodies, with exceptional capacity for precise targeting and binding to pathogens and pharmaceutical targets, have become indispensable tools in cutting-edge research across a wide array of scientific disciplines. So far, nearly 1,200 antibody therapeutics are undergoing clinical studies, with approximately 175 currently in regular review or awaiting approval [1]. Monoclonal antibodies (mAbs) and antibody fragments, which include antigen-binding fragments (Fabs), single chain variable fragments (scFvs), and nanobodies (Nbs), have extensive applications as biological probes and therapeutic agents across diverse fields [2–4]. The advancement of antibodies and antibody-derived macromolecules for various therapeutic objectives underscores the importance of comprehending antibody structure, particularly the intricate relationships between structure and function [1,5,6].

Immunoglobulin (Ig) structure manifests as a Y-shaped glycoprotein comprising two identical heavy chains and two identical light chains (Figure 1) [7]. The amino-terminal variable domains on both the heavy and light chains denoted as V_H_ (for heavy chain) and V_L_ (for light chain), collectively constitute the variable region of the antibody, which allow the antibody to bind to specific antigens selectively [8]. Among the variable domains of the antibody, complementarity-determining region (CDR) loops are the most diverse regions that determine specific antibody binding, contributing to the recognition and binding to the antigen and the vast diversity of antibodies in the immune system [8–10]. CDR-focused affinity maturation and CDR grafting technologies are also often regarded as essential steps for potent therapeutic antibody development [11–13]. These prior studies and technologies further underscore the importance of precisely defining and comprehending the binding mechanisms involving the CDRs of antibodies.

**FIGURE 1.**
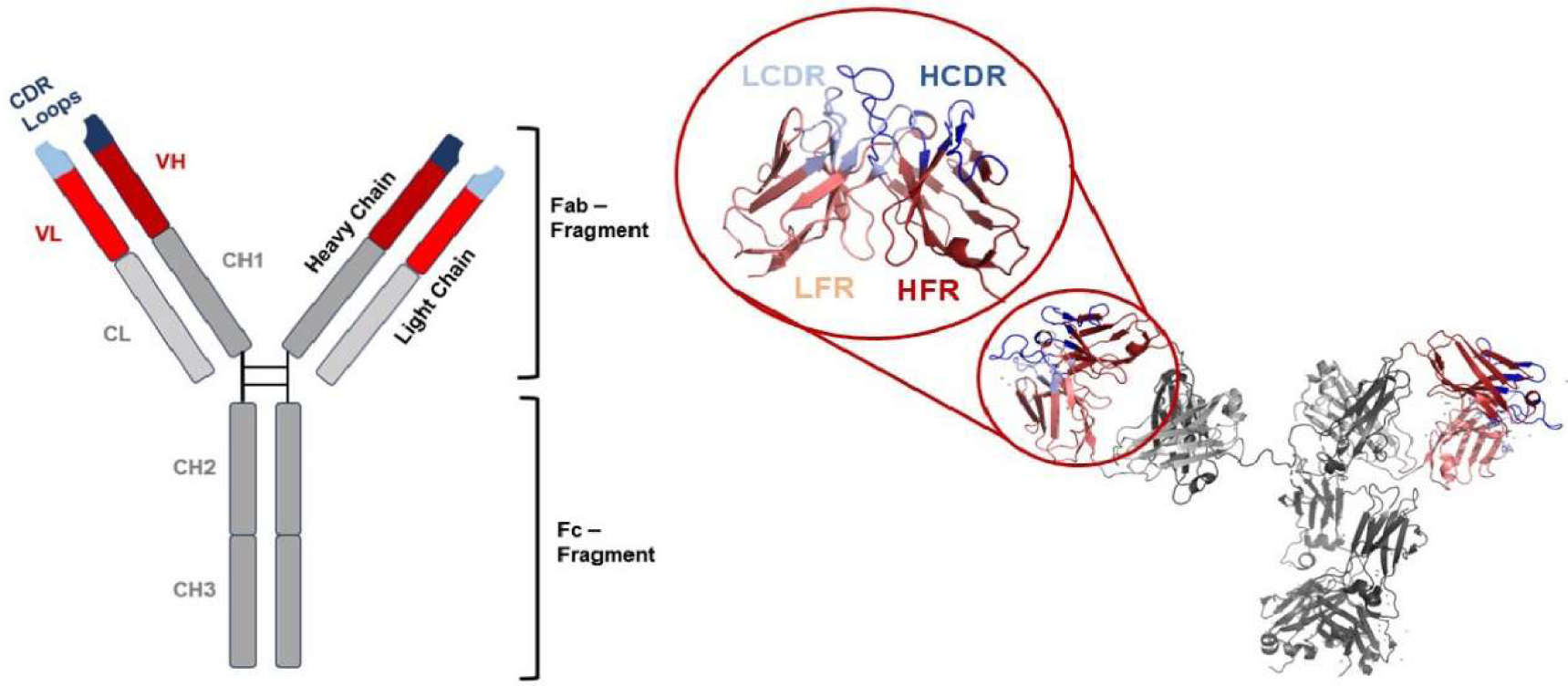
A schematic presentation of the IgG structure (left) and a 3D structure of a human IgG1 (PDB ID: 1HZH, right) [86]. The LCDR, HCDR, Variable light domain (V_L_), Variable heavy domain (V_H_), Constant light domain (C_L_) and Constant heavy domains (C_H_) are color coded in light blue, blue, light red, red, light grey and grey respectively.

To precisely pinpoint the CDR residues, several amino acid numbering systems and CDR definitions have been established. These numbering schemes are founded on the extensive structural and sequence similarities among immunoglobulin frameworks [14]. Antibody numbering schemes allow for effective identification of the CDR residues at their expected positions by aligning the amino acid sequence with these numbering schemes, without solving the structure of each antibody. The first numbering scheme, Kabat, was introduced in 1970 by Kabat and Wu [15]. It was formulated by aligning 77 Bence-Jones proteins and immunoglobulin light chain sequences. This numbering scheme identifies the CDRs based on the residues exhibiting the most diverse sequences [15,16]. The Chothia numbering scheme, pioneered by Chothia and Lesk in 1987, was developed through an in-depth examination of hypervariable domains within high-resolution immunoglobulin structures [17,18]. They identified residues crucial for adopting particular loop conformations, termed “canonical structures,” and the CDRs were identified based on the conservation of structural residues [17]. Note that the earlier Chothia CDR definition encompassed more residues, and in 2021, the Andrew Martin group provided a “consensus CDR” definition on their website [19].

In 2008, Abhinandan and Martin introduced a new numbering scheme known as the Martin numbering scheme, which aimed to rectify the inaccuracies in insertion site assignments in the Kabat numbering scheme [20,21]. The Martin numbering scheme, also known as the “enhanced” Chothia numbering scheme or the “AbM” numbering scheme, is regarded as a more advanced and refined numbering system representing a compromise between the Kabat and Chothia schemes [19,20]. It incorporates multiple corrections and improvements derived from extensive structural analyses and comparisons. In 1997, Lefranc et al. introduced the IMGT numbering system based on amino acid sequence alignment of the germ-line V genes [22]. Later, the IMGT numbering scheme annotation was expanded to encompass the entire variable domain and several useful tools to analyze and visualize the antibody variable sequences based on the IMGT numbering scheme were developed [23]. The IMGT numbering scheme has been regarded as the primary reference in immunogenetics and immuno-informatics [14,24]. The four numbering schemes, Kabat, Chothia, AbM, and IMGT, are widely employed to delineate CDRs, and various online servers like AbRSA and ANARCI, offer alignment tools that facilitate rapid and accurate annotations based on these numbering schemes [25–27]. Several other numbering schemes offer valuable information from other distinct perspectives. The Gelfand numbering scheme, introduced by Gelfand and Kister in 1995, identifies positions within antibody strands and loops that exhibit notable conservation or shared characteristics [28]. In 2001, Honegger and Plückthun introduced Honegger’s numbering scheme (AHo’s), which preserves structural integrity by aligning the Cɑ positions within the most structurally conserved core region, while also highlighting locations suitable for accommodating insertions and deletions [29].

Using diverse amino acid numbering schemes offers insights into antibodies’ structural and sequential aspects, satisfying various research purposes. Despite their significance, there is a notable absence of comprehensive comparisons and assessments among these existing numbering schemes. Also, the current array of available numbering schemes has introduced confusion and the potential for erroneous identification of framework and CDR residues [14]. For instance, a recent investigation into antigen-antibody interactions employed the IMGT numbering scheme for residue identification, while the definition of complementarity-determining region (CDR) residues followed Chothia’s framework, which may contribute to potential confusion in subsequent studies [30]. A study addressing structural aspects presented another instance wherein the combination of the Kabat numbering scheme and the North numbering scheme led to uncertainty during the examination and interpretation of crucial residues within the variable domain of antibodies and complementarity-determining regions (CDRs) [31,32]. A lack of consensus prevails among these various numbering schemes, particularly concerning delineating the boundaries of CDRs. Lastly, compared to the relatively modest database sizes available in the early 2000s, the current landscape boasts an abundance of more than 120,000 variable sequences housed within the abYsis database, while the SabDab database encompasses over 7,700 antibody and antibody fragment structures [33–35]. These large databases underscore the necessity for reevaluating sequential and structural-based numbering schemes in light of the wealth of recent information.

This report presents an extensive analysis of amino acid frequency within immunoglobulin variable domains, aiming to assess the level of sequence conservation among residues designated as CDRs according to various numbering schemes. To evaluate the structural conservation of these CDR residues, we conducted structural alignments across variable domains derived from immunoglobulins from diverse species and subgroups. Furthermore, we performed amino acid sequence alignments between mouse and human immunoglobulin domains to re-examine antibody humanization hotspots. Lastly, we identified and visualized the consensus sequences within the human antibody immunoglobulin variable domain, represented in a 2D map. These findings shed light on regional sequence and structural conservation within antibody sequence databases while also highlighting discrepancies stemming from different numbering schemes. These insights yield valuable guidelines for the precise delineation of antibody CDRs and the strategic design of antibody repertoires, with practical implications in developing innovative antibody-based therapeutics and diagnostics aimed at treating and preventing human diseases.

## RESULTS

### Analysis of immunoglobin data

We analyzed non-identical amino acid sequences derived from an aggregate of 124,317 variable domains of IgGs, sourced from the abYsis database [35]. To evaluate the difference in the distribution of amino acids at each position compared to a reference state, we primarily used relative entropy (RE). RE is an information theoretic measure related to multinomial probability. An RE of zero corresponds to a distribution identical to the reference, while higher RE values signify a greater degree of difference between the measured distribution and the reference state (the maximum depends on the nature of the reference distribution). In the context of sequence conservation level analysis, we defined the reference state as the distribution of amino acids within all of the variable domains in the abYsis database. The possible RE values in this study ranged from 0, representing an identical distribution, to 5, indicating the most divergent distribution.

### Sequence analysis based on the Kabat numbering scheme revealed differing levels of conservation in the CDRs

The residues within CDRs showed a moderately lower level of conservation compared to the framework, but some were more varied than others (Fig. 2A, B and C). The average relative entropy values computed for the entire variable domain, the framework regions, and the CDRs are 1.65, 1.94, and 0.92, respectively (Fig. 2D). (Recall that a relative entropy of zero indicates a frequency distribution that is the same as the reference, where larger values indicate a larger divergence. In general, larger values imply less variation, or conservation.) Among the six individual CDR loops, four have average relative entropy values ranging between 1.0 and 1.2 (Table S1). LCDR3 and HCDR3 exhibited notably lower average relative entropy values of 0.76 and 0.39, respectively, indicating a much higher level of sequence variability within these regions. The median values calculated for each region closely resembled the respective means, except for the HCDR3, where the median was almost 50% lower than the mean. This deviation arises from the notably higher RE values observed at positions H101 and H102 within this region. Both LCDR1 and LCDR3, as well as the HCDR1 and HCDR3, are bordered by two residues (usually a Cys and Trp/Phe pair) that demonstrate a substantially elevated level of conservation.

**FIGURE 2.**
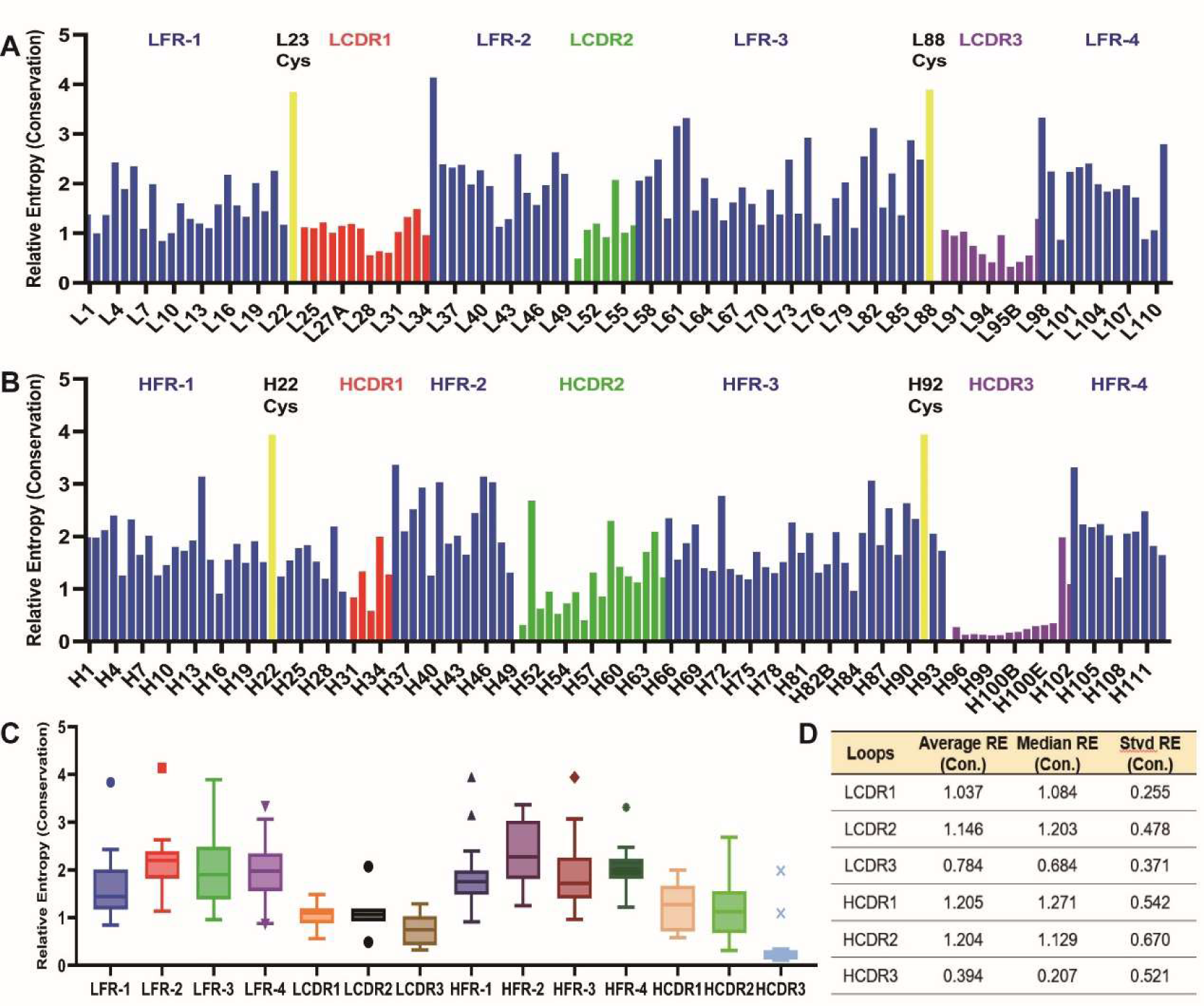
The reside-wise relative entropy calculation based on the Kabat numbering scheme. **A)** RE calculated in the variable light domain with the frequencies of amino acids found within the database of antibody variable domains as the reference state. **B)** RE calculated in the variable heavy domain with the frequencies of amino acids found within the database of antibody variable domains as the reference state. **C)** The box-and-whisker plot of RE values calculated within each region. The boxes of the plots contain data within the interquartile range (IQR), while whiskers spread from the boxes to 1.5 times IQR. The center line in the boxes is the median of the data and points above the whiskers are values which are higher than 1.5 times IQR above the third quartile. n = 23, 15, 32, 13, 30, 14, 22, 11, 14, 7, 11, 5, 17 and 14 for each plot, respectively. **D)** The average (Avg.), median (Mdn.) and Standard deviation (Stvd) of the relative entropy calculated in each loop region.

### Parallel analysis of sequence and structure indicates that Kabat and AbM include more conserved residues in their CDR definition than Chothia and IMGT

We compared the different numbering schemes to the Kabat scheme (Fig. 3A, Table S2). Chothia, AbM, and IMGT generally exhibit a lower RE value in four of the six CDR loops, except for LCDR3 and HCDR1. LCDR2 specifically has a lower average RE in the Chothia and IMGT schemes compared to the Kabat and AbM schemes, which is likely due to the small number of positions in this CDR (usually 2-3) using these two numbering schemes (Fig. S1 and S2). For LCDR3, the average RE values among the four schemes are similar, primarily because the defined range of LCDR3 remains almost identical across these schemes. The exception to this is the Consensus Chothia scheme which excludes L89, L90, and L97, leading to a minor reduction in the average relative entropy (Fig. S1a). Intriguingly, HCDR1 exhibits slightly higher average RE values in the Chothia, AbM, and IMGT schemes, potentially stemming from the Kabat scheme’s incomplete coverage of the entire HCDR1 loop. Meanwhile, in HCDR3, the average RE values is largely consistent in the Kabat, Chothia, and AbM numbering schemes, while the IMGT scheme shows a higher value due to the inclusion of the more conserved residues H105 and H106.

**FIGURE 3.**
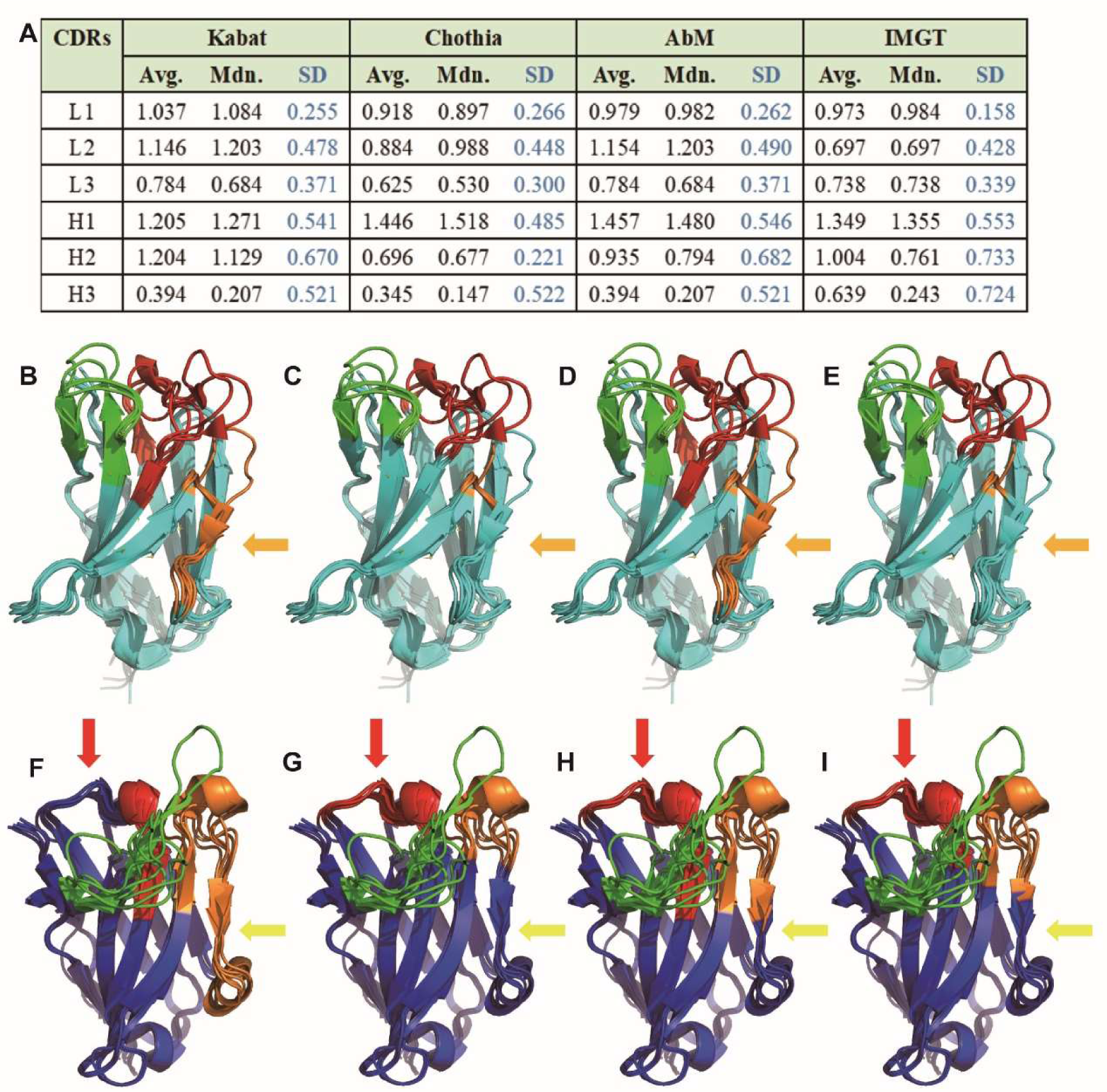
The relative entropy and structural alignments of the CDRs based on different numbering schemes. **A)** The average (Avg.), median (Mdn.) and standard deviation (Stvd) of the RE values calculated based on different numbering schemes. **B - E)** The structural alignment of the 7 selected variable light domain based on the Kabat, Chothia, AbM and IMGT numbering scheme respectively. **F - I)** The structural alignment of the 7 selected variable heavy domain based on the Kabat, Chothia, AbM and IMGT numbering scheme respectively. Color coding: Light Blue: Light chain framework, Blue: Heavy chain framework, Red: LCDR1/HCDR1, Orange: LCDR2/HCDR2, Green: LCDR3/HCDR3.

We mapped the different numbering schemes on to several antibody structures (Tables S3 and S4). While the lambda light chains show significant homology to each other, as do the kappa light chains, there are notable differences between the kappa and lambda light chains (Fig. S3). Several positions in the annotated CDRs exhibit significant structural conservation (Fig. 3B-I). Predominantly, these positions are positioned at the ends of the CDR loops. Moreover, we found two regions of highly conserved structural attributes: one spanning L53 to L56 in LCDR2 and the other encompassing H57 to H65 in HCDR2 (Fig. S4). These regions align with the boundaries designated as CDRs within the Kabat and AbM numbering schemes but not in the Chothia and IMGT numbering schemes (orange arrow in 3B-3E and yellow arrow in 3F-3I). The structural alignment suggests that these two regions do not fall within the loop regions, instead encompassing a conserved beta sheet and an extended loop structure. Some other structurally conserved residues were also identified during structural alignments (Table S5). When these structural findings are compared across different numbering schemes, it becomes apparent that the Kabat and AbM schemes incorporate more structurally conserved residues in their defined CDRs than Chothia and IMGT. It is also noteworthy that both the Chothia and IMGT schemes appear to underestimate the inclusion of residues within LCDR2 when compared to the results obtained from structural alignment (orange arrow in 3B-3E).

Our relative entropy calculations identify several conserved positions with the CDRs as defined by all of the numbering schemes, notably L54 within LCDR2, H29 and H34 within HCDR1, H51 within HCDR2, and H101 within HCDR3 (Fig. S1a and S1b). From the structural alignments, we find that L54 occupies a structurally conserved region positioned at the C-terminus of LCDR2. H29 resides at a pivotal conserved site within HCDR1 that demarcates the division of HCDR1 into two separate loops (see the next section). While H51 and H101 are not structurally conserved according to the alignment, these two residues are situated at the terminal points of the HCDR loops. Specifically, H101 is highly conserved in Asp (frequency 76.9%, RE = 1.98, compared to an average of 0.39 across the entirety of HCDR3).

### A structurally conserved “pivot” point was observed in the CDR1 while showing different preferences of amino acids in the light chain and heavy chain

CDR1 loops have previously been observed to have a distinctive topological characteristic, featuring a structurally conserved residue that falls between two interconnected loops [29,31], but this has received limited attention in sunsequent work. We have also observed a consistently preserved position within most CDR1 loops examined (15 out of 16, Fig. S5). We designate this position as the “pivot” point within the CDR1 loops, situated at L29 and H30 (or H29 in Kabat, AbM & Chothia), which divides the CDR1 into two loops, with its side chain immersed in the hydrophobic pocket between the two beta sheets (Fig. 4A&4C).

**FIGURE 4.**
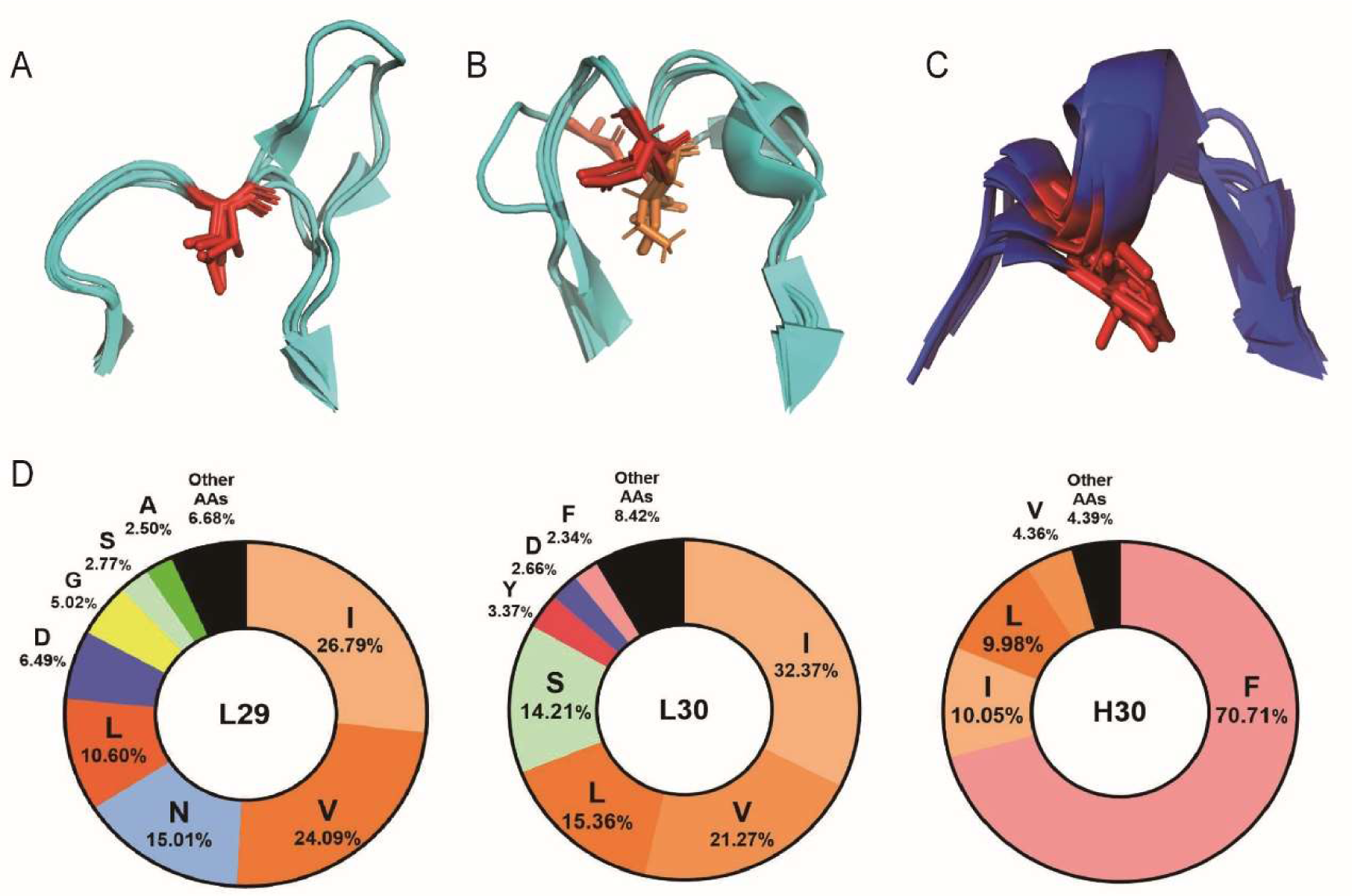
The structure alignment of the LCDR1 and HCDR1, and amino acid distribution at the pivot points. **A)** The structure alignment of the LCDR1 in kappa variable light domain with, L29 shown in sticks and color coded in red. (PDB ID: 2ZKH, 6TCM, 1CR9, 6Z7X, 5VH3) **B)** The structure alignment of the LCDR1 in lambda variable light domain with, L29 and L30 shown in sticks and color coded in red and orange respectively. (PDB ID: 6QBC, 6XRJ, 2YKL, 6H3H) **C)** The structure alignment of the HCDR1 in variable heavy domain with, H30 shown in sticks and color coded in red. (PDB ID: 2ZKH, 6TCM, 1CR9, 6Z7X, 5VH3, 6QBC, 6BE2) **D)** The amino acid distribution at the pivot points based on the IMGT numbering scheme.

In the Kabat numbering scheme, the pivot point is identified as the second inserted amino acid following L27, specifically as L27B. This observation suggests that further modifications to the insertion sites may be required to align with the Kabat scheme. H30 is mostly hydrophobic, primarily driven by a high prevalence of the amino acid phenylalanine, which constitutes 71% of the total residues in this region (Fig. 4D). In contrast, L29 displays a distinct preference for smaller nonpolar residues, with isoleucine, valine, and leucine collectively representing 60% of the overall residue frequency in this position. Phe is a minor constituent at this site, contributing only 0.5%. Furthermore, there is a relatively high occurrence of Asn at this site, in contrast to previous suggestions that pivot points are consistently characterized by high hydrophobicity and their sidechains engage with the hydrophobic core of the domain [29].

Despite the substantial structural disparities observed between kappa and lambda light chain structures, a pivot point is discernible in both types within the LCDR1 region (Fig. 4A-C). In kappa light chains analyzed, this pivot point is at L29 (Fig. S6a). However, in seven out of eight lambda light chain structures examined, the position of the pivot point is shifted to L30, as denoted in the Chothia, AbM, and IMGT numbering schemes (Fig. S6b, Table S6). This shift potentially accounts for the presence of polar amino groups at L29. The only exception to this trend was observed in a lambda light chain derived from an IgG2 variable domain, where no pivot point was evident in the LCDR1 loop [36,37].We also observed a significant preference for small nonpolar amino acids at L30 (IMGT numbering scheme), with Val, Ile and Leu collectively constituting 69% of the total residues in this location (Fig. 4D). Some variations in amino acid distribution occur in the Chothia and AbM numbering schemes due to an inserted point at L30 in these numbering schemes.

Residues located within the first loop of these CDRs, which span from the conserved Cys to the pivot point, consistently exhibited higher relative entropy values compared to residues within the second loop, which extend from the pivot point to the conserved Trp (Table S7a and S7b). This pattern was observed in both LCDR1 and HCDR1. This suggests that the C-terminal loops within the CDR1 regions display greater sequence diversity compared to their N-terminal counterparts. Other studies have also suggested that the C-terminal loops are more intimately involved in direct interactions with antigens in comparison to the N-terminal loops [30,38]. These findings collectively suggest that the C-terminal loops may exhibit more CDR-like characteristics than the N-terminal loops.

### Ser, Tyr, and Gly have increased frequency in the variable domains of immunoglobulin, while Tyr has increased frequency explicitly in the CDRs

We examined the distribution of amino acids in antibodies, compared to the distribution in all proteins in yeast (a neutral reference state, Fig. 5). Compared to proteins overall, the variable domain exhibits an increased frequency of neutral amino acids with small side chain volume, specifically serine, threonine, and alanine. An increase is also seen in the fraction of glycine, tryptophan, and tyrosine.

**FIGURE 5.**
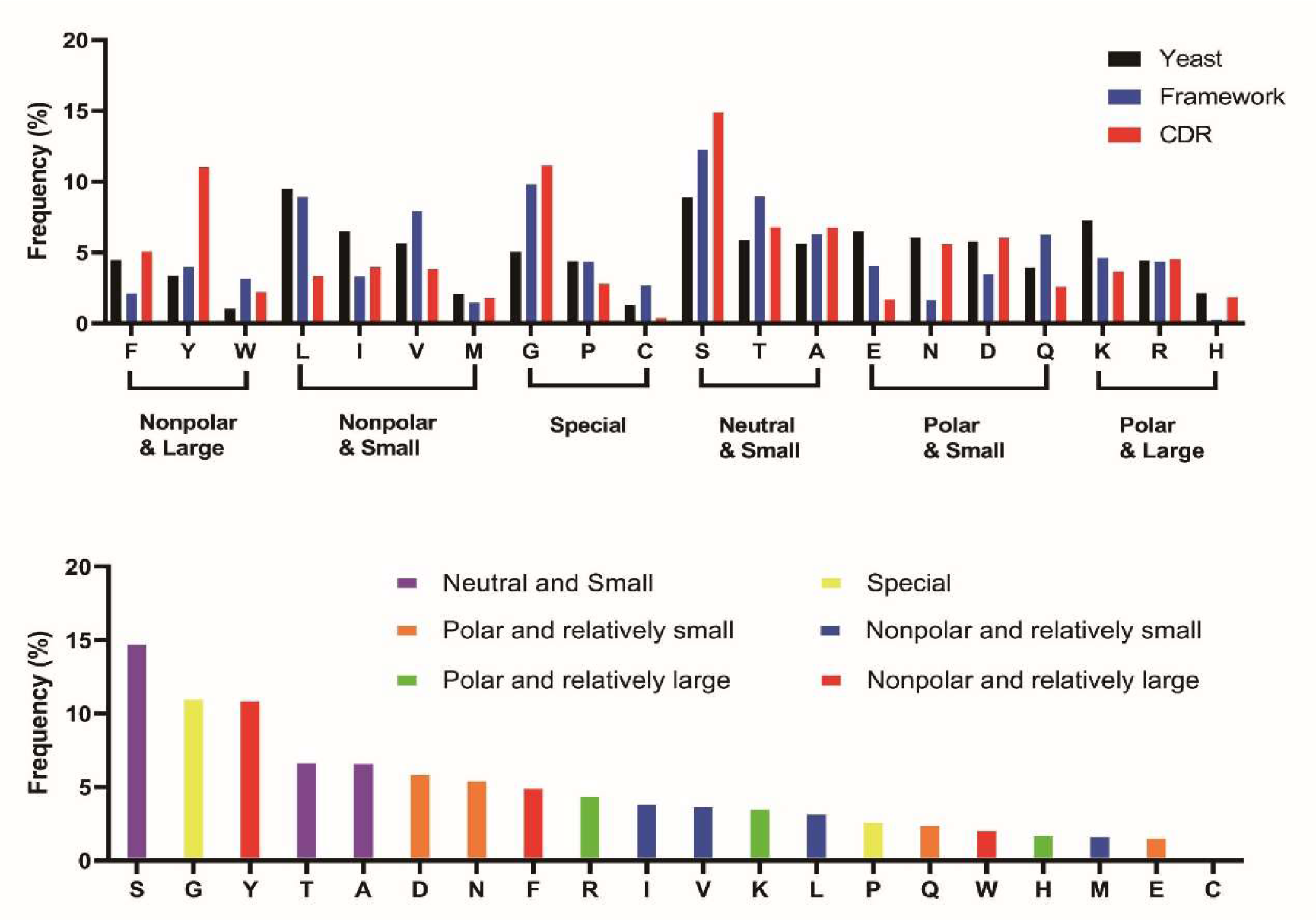
The amino acid distribution in the frameworks and the CDR. The overall amino acid frequency in the framework and CDR regions. The CDRs are defined based on the “general CDRs” (see Results). The reference distribution “yeast” was based on codon usage in the *Saccharomyces cerevisae* genome [68]. **A)** The overall distribution of the amino acids in the framework and CDRs. **B)** The amino acid distribution in the CDRs.

We also examined the distribution of amino acids in the frameworks regions (FRs) and CDRs. Within the FRs, the aggregate frequency of large aromatic amino acids within the variable domain approximately aligns with the proteins overall. However, the FRs, showed a marked increase in the frequency of Gly and Ser within the FRs, with their occurrence heightened by 80% and 50% respectively. Phe is also observed to have lesser prevalence and Trp has an elevated occurrence. Four Trps are highly conserved in the FRs at L35, H36, H47 and H103 in Kabat numbering scheme, with greater than 90% frequency. The beta-branched amino acids Val and Thr also have increased frequency in the beta-sheet structure of the FRs, in agreement with their propensities [39,40].

In the CDRs, a particularly striking higher frequency is seen for Tyr. Previous studies have suggested that the overabundance of Tyr within the CDRs can be attributed to its unique molecular properties [30,41]: the presence of an aromatic ring, combined with the hydroxyl group facilitates the formation of various types of interactions to the antigen, including H-bonds and hydrophobic interactions. Within the CDRs, the percentage of Tyr is 11.0%, compared to the 3.4% observed in proteins overall and the 4.0% in the FRs. Tyr is more common within the HCDRs as well as the LCDR1 & LCDR3 (Fig. S6). But the abundance of Tyr in the LCDR2 is notably lower compared to the other CDR loops. In our CDR amino acid analysis, the frequency of Tyr is lower than that observed in an analysis of paratopes (the residues in contact with the antigen), probably attributed to the broader definition of the consensus CDR employed in our analysis [41]. This observation implies that Tyr is concentrated in regions directly engaging with antigens.

In contrast to previous studies of paratope amino acid frequencies, our CDR amino acid distribution analysis reveals a higher frequency of Ser and Gly [30,41]. Ser is the most prevalent amino acid in the CDRs (14.9%) and is believed to delineate the border of the hydrophobic patches and form H-bonds to the antigens [41]. Gly also emerges as a notable presence within the CDRs, accounting for 11.1%. In contrast, Gly was not enriched in paratope sequences in previous studies [41]. This observation suggests that Gly has a substantial influence on shaping the loop structures but not contacting the antigens. Gly exhibits a significant enrichment in inserted residues, such as H100A, particularly within the HCDR3. This observation also suggests that these amino acids may not be directly involved in antigen interaction but likely serve auxiliary roles in maintaining loop structures and interacting with solvent molecules.

### Sequence analysis between human and mouse variable domains refined several “hot spots” for antibody humanization

The process of antibody humanization is a method of transforming monoclonal antibodies (mAbs) obtained through the immunization of mice or other animals into sequences that closely resemble human antibodies. This conversion is undertaken to alleviate any undesirable clinical characteristics, like the development of human anti-mouse antibodies (HAMA) [42].

To humanize potential clinical antibodies, various strategies have been developed to substitute the original sequence, to eliminate immune response against the murine antibody while preserving its specificity, affinity, and stability. The most traditional approach is by CDR-grafting and back mutations [43]. CDR-grafting involves selecting the CDR loops from a given murine sequence and integrating them into the human framework region that exhibits the highest homology to the original framework sequences [43,44]. The objective is to create a molecule that maintains stability and activity but enhances tolerance by the human immune system by combining human frameworks with the original murine CDR [44]. Some additional strategies have been developed to guide the selection of the grafted CDR residues and the homologous framework. Specificity-determining residue (SDR) grafting reduces the potential immunogenicity by minimizing the region that has been grafted to the human antibody template [45]. Several computational programs, such as CoDAH and CUMAb, were developed to guide amino acid substitution in the humanization process by considering both the evaluation of humanness and structural stability criteria [46–48]. More recently, a deep-learning-based platform called BioPhi has been developed, offering a novel method for humanization through deep learning trained on the observed antibody space using language modeling. This approach has demonstrated promising results in silicon humanization [49].

A few humanization approaches of antibody variable domains were developed under the guidance of the statistical analysis of the human germline [44,46,48,50–52]. The basic logic of these approaches is to maximize the similarity to the closest human germline sequences and a term was introduced to evaluate the similarity is called “humanness score” [44]. The humanness score has slightly different definitions among different approaches, including the average similarities of the sequences, average similarity among the closest 20 sequences or the similarity of the immunogenicity relevant residues [48,50,51]. Further studies optimize the approaches by incorporating the factor of pair correlations between residues. More recently a deep-learning based computational approach has been developed to evaluate the humanness of sequences based on the likelihood of the given sequence belonging to the distribution of native variable domain (Fv) sequences derived from human, called AbNatiV [53]. Though these evaluation approaches provide us a fast and thorough screening for the potential residues that are not similar to human germline in an antiobdy of interest, it is beneficial for us to identify the residues compositions that are different among mouse and human species on a broader scale of statistical analysis. The up-to-date statistical analysis of the sequences can also further provides guidance for mutagenesis after figuring out the potential sites for humanization. Furthermore, despite the availability of multiple computational strategies, the humanization process still heavily relies on trial-and-error, which can be benefit from our statistical analysis [44].

We conducted an analysis of the amino acid distribution within the variable domains of both human and mouse antibodies. The amino acid distributions between these groups are highly similar, but some differences are evident: mouse antibodies have a slightly lower occurrence of Gly and a slightly higher prevalence of polar amino acids within their sequences compared to human antibodies (Fig. S7). Specifically, Val, Gly, and Arg are less common. In contrast, Tyr and Lys are more common.

We examined the sequence divergence between human and mouse antibodies (Fig. 6 and S8). The average RE values within the framework regions of both the variable heavy and variable light domains are similar (Table S8a and S8b). The LCDRs, on average, show a higher average RE compared to the HCDRs. This finding suggests that a greater level of differences between mouse and human sequences exists within the LCDRs. In the framework regions, Light Framework Region 1 (LFR-1) and Heavy Framework Region 3 (HFR-3) exhibit higher average and median RE values compared to the other framework regions. Conversely, Heavy Framework Region 4 (HFR-4) displays relatively lower RE values, with only 2 out of its 11 residues (H108 and H109) exhibiting higher RE values than the average RE value of the heavy chain frameworks.

**FIGURE 6.**
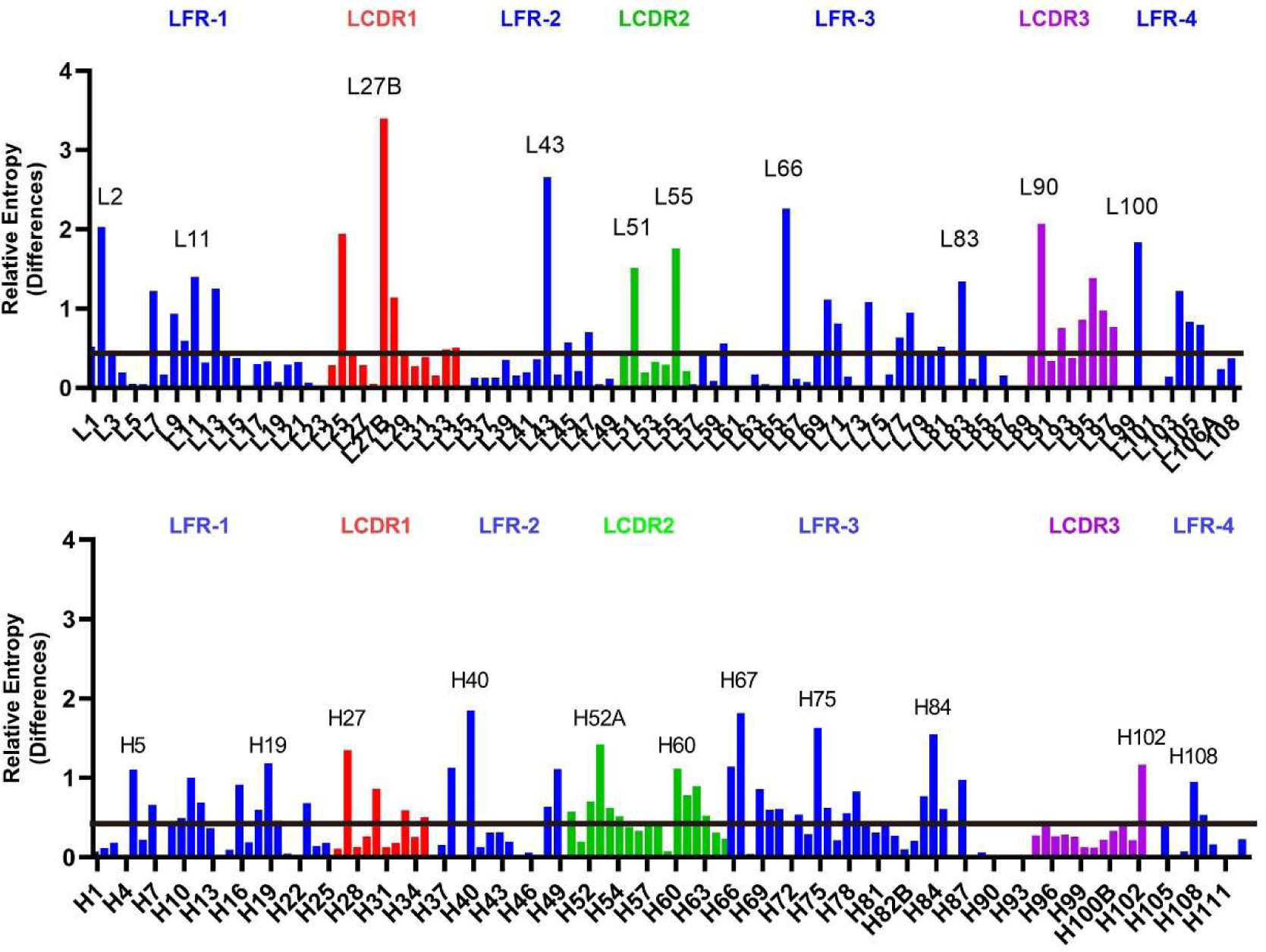
Comparison of human antibody positional distributions to mouse antibodies. The RE values were calculated for each residue in the CDRs and the frameworks using the amino acid distributions in each position in mouse antibodies as a reference state. The CDRs are defined based on the Kabat numbering scheme. The CDRs were color coded in red (CDR1), green (CDR2) and purple (CDR3). The residues with higher RE (higher than the average plus 1.65 times the standard deviation of the region) were marked out with residue numbers. The global average RE value was marked as a solid line.

L27B registers the highest RE value (3.394) among all the residues examined in our study (Table S9). Notably, L27B coincides with the “pivot” point discussed above. Conversely, the pivot point in the heavy chain (H29), Phe dominates the residue composition, and no discernible differences were observed between the mouse and human variable domains (Fig. S9). At the pivot site in the LCDR1, L27B exhibits a higher frequency of polar residues in human immunoglobulins, with 45% consisting of Asn and 19% of Asp. In contrast, the mouse variable domain at L27B has a predominance of nonpolar residues, with negligible or minimal presence of Asn or Asp (<1%). We also conducted a parallel analysis of the equivalent residue in the IMGT numbering scheme, denoted as L29, where similar distribution patterns were observed. In the IMGT scheme, lower yet still substantial amounts of Asn (20%) and Asp (9%) were observed in the human variable domain, while minimal to no Asn or Asp was detected at L29 in the mouse variable domains. The presence of polar amino acids at L29 likely arises due to the influence of the lambda light chain. The divergence in polar amino acid distribution of human and mouse antibody at this point can be attributed to the relatively rare occurrence (∼5%) of the lambda light chain in mice [54–56]. In contrast, the average κ to λ light chain ratio is 2:1 in human [56]. This observation further supports our hypothesis of pivot point distinctions between lambda and kappa light chains. Also, the disparity of the lambda light chain in the human and mouse antibodies may explain the higher levels of differences in the light chains compared to the heavy chains.

To further validate our findings of the hotspots for the humanization on the variable domain based on the relative entropy calculation, we analyzed the consensus mouse antibody variable domain sequences derived from the abYsis database, using the AbNatiV servers and compared the outcome with our RE results (Fig. S10). Seventeen of the 21 humanization hotspots we identified in the RE analysis showed lower AbNatiV scores or had the same amino acid with the human consensus sequence (Table S10). The other 4 residues we identified were either in CDRs (L27B, L51 and H27), or the consensus mouse sequence input was the second most abundant at this residue in human database, which might cause the AbNatiV to recognize them as human sequences (H27 and H75). Conversely, the residues with low AbNatiV scores showed a higher RE value (0.806) compared to the overall average RE value of the variable domain (0.475) with the distribution of human amino acids at each position in the database as the reference state. The results from the AbNativ server also validate our findings of LFR1 and HFR3 having the most residues that are different between mouse and human antibody sequences. Overall, the results from the AbNatiV server are consonant with our RE analysis.

### Consensus amino acid frequency analysis reveals regional conservation in the variable domains of human immunoglobulin

To help visualize the relationship between sequence and structure conservation, a 2D consensus sequence map was generated based on the Honegger numbering scheme, and the beta strands in the beta sheets were classified, highlighted and labeled based on the Gelfand numbering scheme (Fig 7A and B). Most conserved (high RE) residues are situated within regions that are also structurally conserved, underscoring the close alignment between these two dimensions of conservation. Also, when comparing the conservation of the “inner” beta sheet, which interacts with the “inner” beta sheet in the variable light domain, with that of the “outer” beta sheet, the “inner” beta sheet demonstrates a markedly higher degree of sequence conservation. However, this phenomenon is not mirrored within the light chain.

**FIGURE 7.**
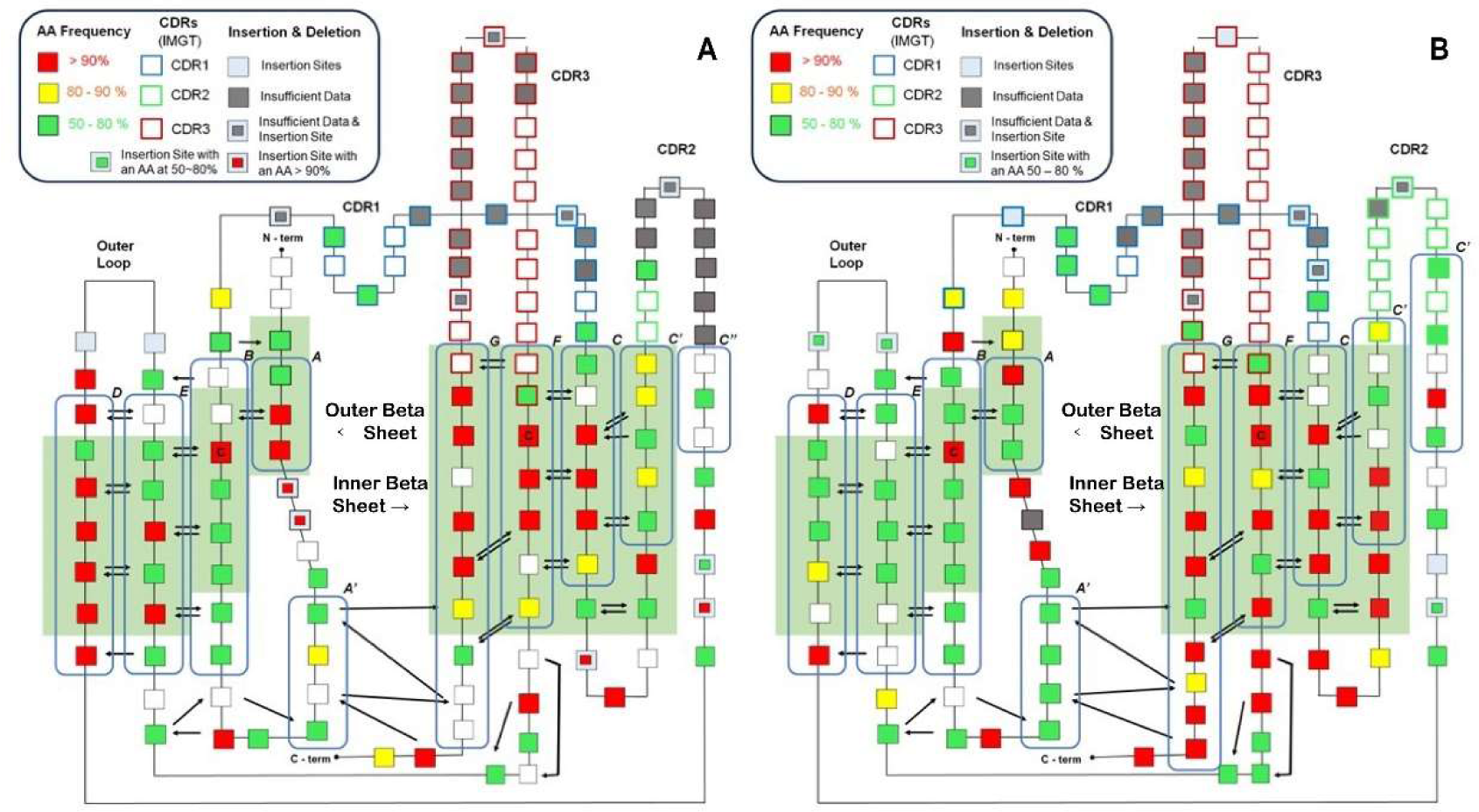
A representative of amino acid spatial distribution and their conservation level of homo sapiens consensus sequences. Each square in this figure represents a residue and the color code corresponds to the relative frequency of the most frequent amino acid at this point. The sequence was numbered using the Aho numbering scheme and the CDR were marked using the IMGT numbering scheme. The arrows between residues represent the hydrogen bonds between the backbones. The green solid square in the background corresponds to C_α_ conserved region defined by the Aho numbering scheme and the blue frames with a letter on the top right corner represent the beta strands and their title based on the Gelfand numbering scheme. The highly conserved Cysteines (C23, C106 in both domains) were marked with a “C” in the box. **A)** The 2D map of the variable light domain **B)** The 2D map of the variable heavy domain

There is also a remarkably conserved beta strand within the variable light domain. This beta strand is defined by the “RFSGSXSG” pattern and spans positions L75 to L82 according to the IMGT numbering scheme (Fig. S10a and S10b). The conservation for these residues exceeds 90%, with the exception of L80, in which Gly is observed at 58%. This particular beta strand is solvent-exposed, yet its function remains enigmatic, as no definitive explanation for the purpose of this highly conserved pattern has been identified to our knowledge.

Also, several positions in the HCDR3 (based on IMGT definition), have very low variation. Specifically, at position H105, Ala is present at a frequency of 85%, while Arg occurs at 70% at position H106 and Asp is seen at 74% at position H116. These highly conserved residues were also observed in previous studies, and are found in both human and mouse antibody sequences [30,57]. H105 and H106 were only defined as HCDR3 based on the IMGT numbering scheme but not in Kabat, Chothia and AbM, which explains the relatively higher RE of the HCDR3 based on the IMGT numbering scheme compared to the others. The conservation of these residues is likely due to the rearrangement of the V(D)J genes. In the IMGT numbering scheme, the HCDR3 includes the rearranged Ig V genes (105-106) and V-D-J genes (107-117) [23]. The Ala and Arg showed at 105 and 106 positions are highly conserved in the IGHV germlines in the previous study [58]. The conservation of the Asp at H116 is likely due to the high prevalence of the Asp at this position in all of the IGHJ genes except for the low frequency IGHJ1 genes (<4%) [59].

## DISCUSSION

Numbering schemes for antibodies offer a useful way to identify CDRs in antibody sequences, using various methods. However, the widespread adoption of new schemes is hindered by discrepancies between different numbering schemes and a lack of comprehensive evaluation, especially given the wealth of data available in today’s databases [14]. Utilizing statistical analysis to investigate residue-wise conservation across 124,317 amino acid sequences within the variable domains found in the abYsis database, our findings indicate that the Kabat and AbM numbering schemes tend to encompass more structurally and sequence conserved residues when compared to the IMGT and Consensus Chothia numbering schemes. Contrastingly, the definitions of CDRs by IMGT and Consensus Chothia seem to be more rigorous, incorporating only the most diverse loop residues. However, they also omit several residues in LCDR2.

The results of this study show that even with the massive database of antibody sequences today, the CDRs can still be identified by their diversity compared to the antibody frameworks. HCDR3 exhibits the highest diversity, consistent with previous knowledge, while the substantial diversity observed in LCDR3 has been underrepresented in prior research. Sequence and structural alignments reveal that the major numbering schemes (Kabat, Chothia, AbM, and IMGT) exhibit similar CDR definitions for LCDR1, LCDR3, and HCDR3 with minor distinctions. However, discrepancies arise in LCDR2 and HCDR2, where Kabat and AbM overestimate CDR boundaries, while Chothia and IMGT underestimate them. Notably, Kabat’s HCDR1 definition is incomplete and incorrectly positions the insertion point in the N-terminus loop rather than the C-terminus loop.

The analysis of amino acid frequencies began with an examination of HCDR3, revealing that the top 5 amino acids predominantly used in HCDR3 were “YGTSA” in naïve antibodies and “YGSDA” in synthetic antibodies [60,61]. Our findings align with their results, and in the case of the top 6 amino acids identified in the CDR loops in our analysis are “SGYTAD.” The different rank of the most frequent amino acid is likely due to the fact that the general CDR definition we employed encompasses residues that were not classified as CDRs in the IMGT numbering scheme used in their analysis. And the amino acid exhibiting the most substantial increase in CDRs compared to the framework in our analysis, Tyr, was also found to have high occupancy in the paratope structure analysis, indicating its crucial role in mediating interactions with antigens [30,41]. However, the Gly and Se, which exhibited high frequency in our CDR analysis, did not exhibit distinguishable patterns in the paratope analysis [30,41,41]. The high occurrence of Gly and Ser, as observed in prior studies, can be attributed to Gly contributing conformational flexibility and Ser, with its small and neutral side chain, playing a valuable role in molecular recognition. [61,62].

We also observed a structurally conserved residue within the CDR1 loops that possesses a distinctive pivotal structural characteristic, referred to as the “pivot point” in this study. This residue exhibits a strong preference for hydrophobic amino acids, with its side chain deeply embedded within the hydrophobic core of the variable domain. Nonetheless, the annotation of this pivot point uncovers a minor discrepancy common to all four prominent numbering schemes (Kabat, Chothia, AbM, and IMGT). This discrepancy pertains to the different numbering of this topological equivalent pivot point in kappa and lambda light chains, resulting in variations in amino acid frequencies. Our results highlight the necessity for modifying the current CDR numbering schemes and suggests that researchers should select the numbering schemes wisely based on different research purposes.

Over the past several decades, numerous alterations have been implemented to adjust the existing numbering scheme and rectify a number of discrepancies resulting from the limited database size initially available [14,17,19,20].The concept of pivot points (L29 and H30), as discussed in this study, was initially introduced by Honneger and Plückthun during the introduction of the aHo numbering scheme [29]. In their scheme, they described this pivot point as a typically hydrophobic residue occupying a crucial position that divides CDR1 into two distinct loops [29]. However, the examination of the pivot point concept did not progress further in the development subsequent numbering schemes, and CDR1 was consistently considered a single unit rather than two connected loops. In the Chothia, AbM, and Aho numbering schemes, the insertion sites in LCDR1 were consistently positioned exclusively within the C-terminus loop, indicating a greater degree of length variation compared to the N-terminus loop [17,20,29]. Recent structural analysis has revealed that the C-terminus loop displays increased diversity in amino acid sequences and engages in more extensive interactions with protein antigens. This underscores the importance of separately addressing these two loops in further discussions [30]. Additionally, a comparison of mouse and human antibody sequences highlights the most significant variation at L29, precisely where the pivot point is located in the light chain, likely due to differences in the ratio of lambda and kappa light chains between species.

Our studies of amino acid frequency and relative entropy, utilizing various numbering schemes, offer an evaluation of the existing numbering scheme in terms of sequence conservation and diversity. Subsequent structural alignments and assessments further unveil several discrepancies within these numbering schemes. These findings can aid researchers in selecting the most suitable numbering scheme for their specific research objectives. These insights provide valuable guidance for accurately defining antibody CDRs and strategically designing antibody repertoires, offering practical implications for the development of innovative antibody-based therapeutics and diagnostics to address and prevent human diseases.

## MATERIALS AND METHODS

### The Ab Variable Domain Sequence Data Set

The variable domain sequences of antibodies (referred to as Ab variable domains) employed in this study were obtained from the abYsis database in July, 2023 (http://www.abysis.org/). Our investigation encompassed non-identical sequences derived from a total of 36,479 variable light domains and 87,838 variable heavy domains [35]. In the context of analyzing variations across different organisms, we processed a dataset comprising 48,402 variable heavy domains and 20,197 variable light domains sourced from antibodies inherent to *Homo sapiens*. Concurrently, for the murine investigation, an examination was conducted on a set of 7,325 variable light domains and 23,303 variable heavy domains. The antibody CDRs have been annotated with abYsis using the Kabat, Chothia, AbM (Martin), Honnegger (Aho) and IMGT numbering schemes. The annotation of Gelfand numbering scheme was generated by manual alignment based on the published methods ([28,29]). Positions with less than 30% occupancy have been discarded during the analysis.

### Comparison of CDR definition and insertion sites for different numbering schemes

The alignment of the multiple numbering schemes based on the Kabat, Chothia, AbM and IMGT methods was performed as previously described [15,17,19,20,23]. In this study, we employed an updated consensus Chothia CDR definition, which was formulated by the Martin group in 2021 on their group page (http://www.bioinf.org.uk/abs/info.html) [19]. This revised definition is a culmination of Chothia CDR definitions from previous literature, and the consensus Chothia numbering scheme annotates fewer residues as CDR compared to the original scheme [17,18,63].

To accommodate the variability in length observed in different antibody frameworks and CDR segments, several “insertion sites” have been introduced to ensure that conserved residues share similar numbers in the different schemes. The majority of these insertion sites were strategically placed within CDR loops, where higher levels of length variation are typically observed (e.g., L27A) [64]. Additional insertion sites have been incorporated into the framework to account for variations in length across antibody scaffolds or species, such as the introduction of L106A to accommodate extra amino acids in the lambda light chain compared to the kappa light chain [29,65]. In numbering schemes such as Kabat, Chothia, and AbM, these insertion sites are annotated with letters followed by residue numbers (e.g., L27A, L27B). The IMGT numbering scheme differs in its approach, counting residues continuously from 1 to 128 based on the germ-line V sequence alignment. Consequently, only a few insertion sites are utilized at specific positions (e.g., 111.1-112.1 in HCDR3) and no number is assigned when a residue is absent in a particular sequence. For Instance, in a 8-amino acid-long CDR1 using IMGT, residue 29 is directly followed by residue 34, and residue numbers 30–33 are omitted [14,22–24].

In this work, to analyze the CDR amino acid distribution and the compare the differences between human and mouse amino acid distributions., we introduced a universal definition of the CDR. This comprehensive definition, referred to as the “general CDR,” encompasses all residues defined as CDRs by any numbering scheme, ensuring a consistent and inclusive approach. The definition of CDRs according to different numbering schemes is listed in Table 1.

**TABLE 1.** Comparison of CDR numbering schemes of variable regions defined using Kabat, Chothia, Consensus Chothia, AbM and IMGT methods. General CDRs, defined to include all residues defined as CDRs by any numbering scheme. The general CDR residues are marked following the Kabat numbering scheme. Note that the Chothia CDR discussed later is using the “consensus” Chothia numbering scheme. The insertions sites were identified and adapted from abYsis database [35]

| Loop | Kabat | Chothia | Consensus Chothia | AbM | IMGT | General |
| --- | --- | --- | --- | --- | --- | --- |
| LCDR1 | L24— L34 | L24— L34 | L26— L32 | L24— L34 | 27— 38 | L24— L34 |
| LCDR2 | L50— L56 | L50— L56 | L50— L52 | L50— L56 | 56— 65 | L50 – L56 |
| LCDR3 | L89— L97 | L89— L97 | L91— L96 | L89— L97 | 105— 117 | L89 – L97 |
| HCDR1 | H31— H35 | H26— H32 | H26— H32 | H26— H35 | 27— 38 | H26 – H35 |
| HCDR2 | H50— H65 | H52— H56 | H52— H56 | H50 – H58 | 56— 65 | H50— H65 |
| HCDR3 | H95— H102 | H95— H102 | H96— H101 | H95— H102 | 105— 117 | H93 – H102 |
| Insertion Sites | L27A— L27F,<br>L39A, | L30A – L30F,<br>L39A, | L30A – L30F,<br>L39A, | L30A – L30F,<br>L40A, | 30 – 35 (L),<br>46.1 (L), |  |
|  | L54A – L54E, | L54A – L54E, | L54A – L54E, | L52A – L52E, | 57 – 64 (L), |  |
|  | L66A – L66H, | L66A – L66H, | L66A – L66H, | L68A – L68H, | 84.1— 84.6 (L) |  |
|  | L95A – L95F, | L95A – L95F, | L95A – L95F, | L95A – L95F, | 111.1-112.1 (L), |  |
|  | L106A, | L106A, | L106A, | L106A, | 127 (L), |  |
|  | H6A – H6D, | H6A – H6D, | H6A – H6D, | H7A – H7D, | 10 – 10.3 (H), |  |
|  | H35A – H35H, | H31A – H31H, | H31A – H31H, | H31A – H31H, | 32 – 32.4 (H), |  |
|  | H52A – H52G, | H52A – H52G, | H52A – H52G, | H52A – H52G, | 60.1 – 60.4 (H), |  |
|  | H82A – H82E, | H82A – H82E, | H82A – H82E, | H72A – H72E, | 84 – 84.2 (H), |  |
|  | H100A – H100T. | H100A – H100T. | H100A – H100T. | H100A – H100T. | 111.1-112.1 (H), |  |

### Relative Entropy Analysis

In this work, the degree of conservation pertaining to individual positions has been assessed using relative entropy (RE), which is at times denoted as “D.” Relative entropy is an information-theoretic construct which quantifies the disparity between a given distribution and a reference distribution. This metric is logarithmically related to the multinomial probability associated with observing a specific distribution from a population with the reference distribution. For instance, the analysis of positional sequence conservation, can be found in our previous work [66,67].

The reference distribution used here to compute conservation levels was based on the frequencies of amino acids found within the database of antibody variable domains. This selection offers a reliable approximation of the expected prevalence of amino acids within a standard immunoglobulin variable domain, mitigating the bias introduced by amino acid preferences inherent in such domains (Fig. 8). For comparison, the amino acid usage in all ORFs in *Saccharomyces cerevisae* is shown [68].

**FIGURE 8.**
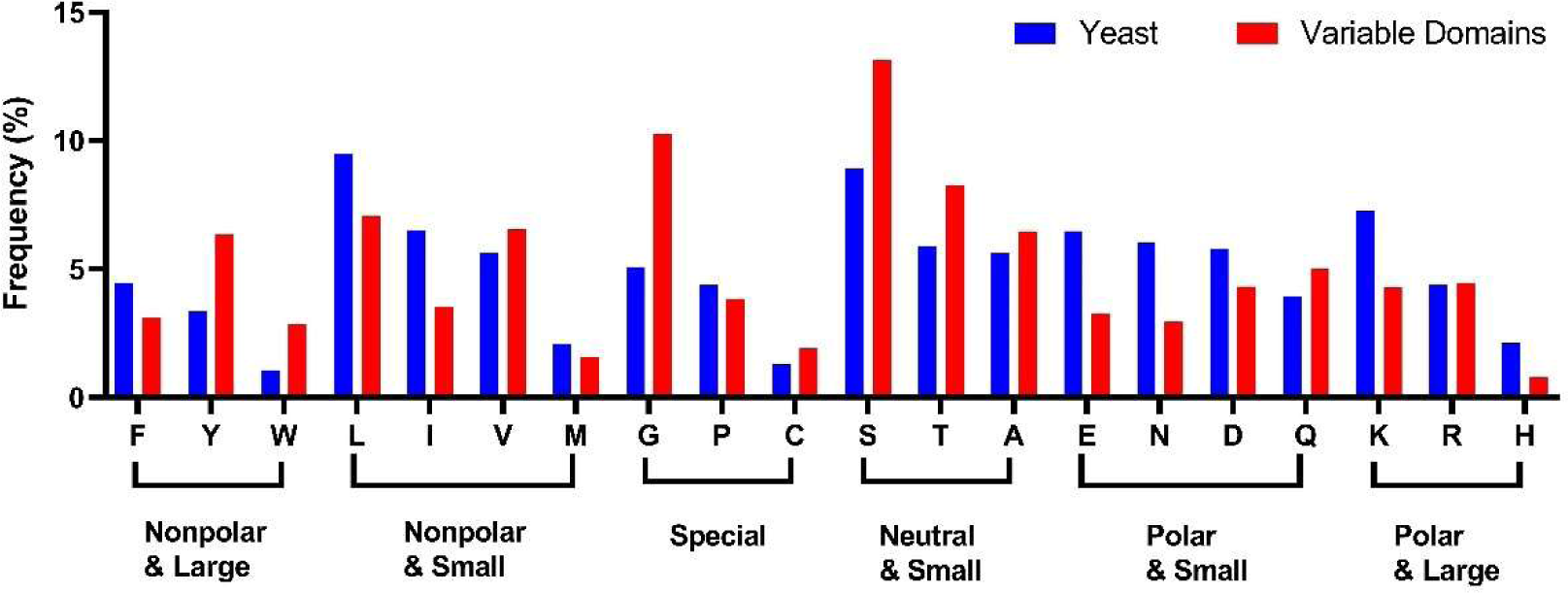
The amino acid distribution in immunoglobulin variable domain. The distribution is used as the neutral reference state, compared to the codon usage in yeast [68].

In cases where a position exhibit pronounced conservation, its amino acid distribution would substantially deviate from the reference, consequently yielding an elevated relative entropy value in relation to a neutral reference state. The RE is calculated as

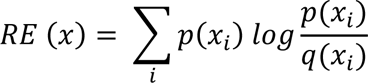

where *p*(*x_i_*) is the frequency of observing residue *i* at position *x* in the sequence alignment and *q*(*x_i_*) is the frequency of observing residue *i* in the reference (Table S10).

To evaluate hot spots for humanization, we quantified the dissimilarity, or “distance” between the amino acid distribution observed at each position in *Homo sapiens* compared to the distribution in *Mus musculus*. In this assessment, the amino acid frequencies at each specific position within the antibody variable domain sequences of both *Homo sapiens* and *Mus musculus* were calculated separately. In this case, the frequencies of amino acids in mouse were used as the reference distribution to calculate the relative entropy.

### AbNatiV Score Calculation

The AbNatiV scored were obtained from the free online server AbNatiV provided by the Department of Health, University of Cambridge (https://wwwcohsoftware.ch.cam.ac.uk/index.php/abnativ) [53].

### Structural Alignment

The structural information and amino acid sequences of seven antigen-binding fragments (Fabs) were sourced from the Protein Data Bank (PDB ID: 2ZKH, 6TCM, 6Z7X, 5VH3, 1CR9, 5L88, 5OPY. 4M43, 6QBC, 4YNY, 5OD0, 6XRJ, 2YKL, 6H3H, 1MFD, 6BE2) [36,69–84]. The CDRs defined according to different numbering schemes (Kabat, Chothia, Martin, IMGT and general CDR) were numbered using the abYsis server and annotated based on Table 1. The predicted CDRs delineated by the Chothia numbering scheme were subsequently refined using the “consensus” Chothia numbering scheme. To facilitate the global structural alignment, the PyMOL software (Pymol^TM^ 2.5.5) was employed, with a focus on displaying only the variable domain or the CDR loop of interest [85].

## ACKNOWLEDGMENTS

KSO, an undergraduate student at the College of Wooster, was supported by a summer internship from the Ohio Five-OSU Summer Research Undergraduate Experience during this work.

## ABBREVIATIONS

CDR: Complementarity Determining Region
scFv: single-chain variable fragment
LCDR: Complementarity Determining Region on the variable light domain
HCDR: Complementarity Determining Region on the variable light domain
IQR: interquartile range
V_L_: variable heavy domain
V_H_: variable heavy domain
C_L_: constant light domain
C_H_: constant heavy domain
mAbs: monoclonal antibodies
Fab: antibody binding fragments
Nbs: nanobodies
Ig: immunoglobulins
AbM: The Martin numbering scheme
Aho: The Honneger’s numbering scheme
RE: Relative Entropy
HAMA: Human Anti-mouse Antibodies.

## APPENDIX A. SUPPLEMENTARY DATA

Supplemental Figure S1-1

Supplemental Table 1-11

## AUTHOR CONTRIBUTION

**ZZ:** Conceptualization, data curation, formal analysis, investigation, visualization, methodology, writing original draft, writing review and editing. **KSO:** Data curation, formal analysis, investigation, writing review and editing. **TJM:** Formal analysis, investigation, methodology, project administration, writing review and editing, supervision, funding acquisition, resources.

