## Supplemental Files for "50 years of antibody numbering schemes: a statistical and structural evaluation reveals key differences and limitations"

Items included in this document:

1) Supplementary Figure 1a: Relative entropy of conservation in the variable light domain

2) Supplementary Figure 1b: Relative entropy of conservation in the variable heavy domain

3) Supplementary Figure 2a: Length of the LCDRs based on different numbering schemes

4) Supplementary Figure 2b: Length of the HCDRs based on different numbering schemes

5) Supplementary Figure 3: Structural comparison between the lambda and kappa light chain

6) Supplementary Figure 4: Structural alignment of the CDR loops

7) Supplementary Figure 5a: Pivot point structures in lambda light chain

8) Supplementary Figure 5b: Pivot point structures in kappa light chain

9) Supplementary Figure 6: Amino acid distribution in each of the CDR loops

10) Supplementary Figure 7: Amino acid distribution in mouse and human antibody

11) Supplementary Figure 8: The whisker plot of the regional relative entropy differences

12) Supplementary Figure 9: Amino acid distribution at the pivot points in mouse and human

13) Supplementary Figure 10: The Relative Entropy and the AbNatiV Score of variable domains

14) Supplementary Figure 11a: 2D map of consensus sequence of human variable light domains

15) Supplementary Figure 11b: 2D map of consensus sequence of human variable light domains

16) Supplementary Table 1: Relative entropy calculated in regions in the variable domains

17) Supplementary Table 2: Comparison of the CDR definition of different numbering schemes

18) Supplementary Table 3a: Antibody template PDB entries, organism, reference information

19) Supplementary Table 3b: Antibody template amino acid sequences

20) Supplementary Table 4a: Antibody Template LCDRs amino acid Sequences

21) Supplementary Table 4b: Antibody Template HCDRs amino acid Sequences

22) Supplementary Table 5: Sequential and structural conserved residues identified in CDRs

23) Supplementary Table 6: Lambda and kappa light chain template amino acid Sequences

24) Supplementary Table 7a: Relative Entropy of the Residues in LCDR1

25) Supplementary Table 7b: Relative Entropy of the Residues in HCDR1

26) Supplementary Table 8a: Average relative entropy of different species in each region

27) Supplementary Table 8b: Statistics of the RE of different species and outliers in each Regions

28) Supplementary Table 9: Amino acid distributions at the diverse residues in human and mouse

29) Supplementary Table 10: The RE value and the AbNatiV Score of variable domain residues

30) Supplementary Table 11: Amino acid relative frequency in variable domains

**Figure S1a**

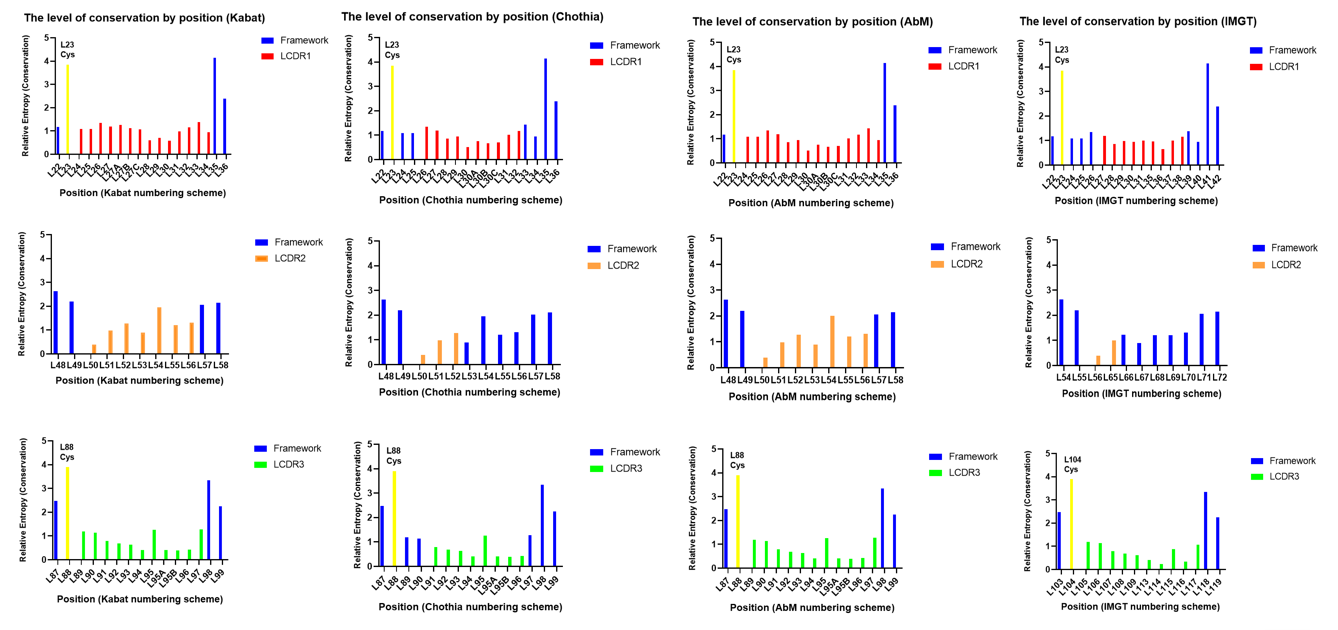
**Supplemental Figure 1a. Relative entropies of conservation for selected residues in the variable light domain.** The relative entropy of conservation was calculated for each residue, with residue position given by the Kabat, Chothia, AbM and IMGT numbering schemes. Relative entropies of residues within the CDR are colored according to the LCDR to which they belong (LCDR1 shown in red, LCDR2 in orange, and LCDR3 in green), while framework residues are shown in blue. Highlighted cysteine residues are shown in yellow.

**Figure S1b**

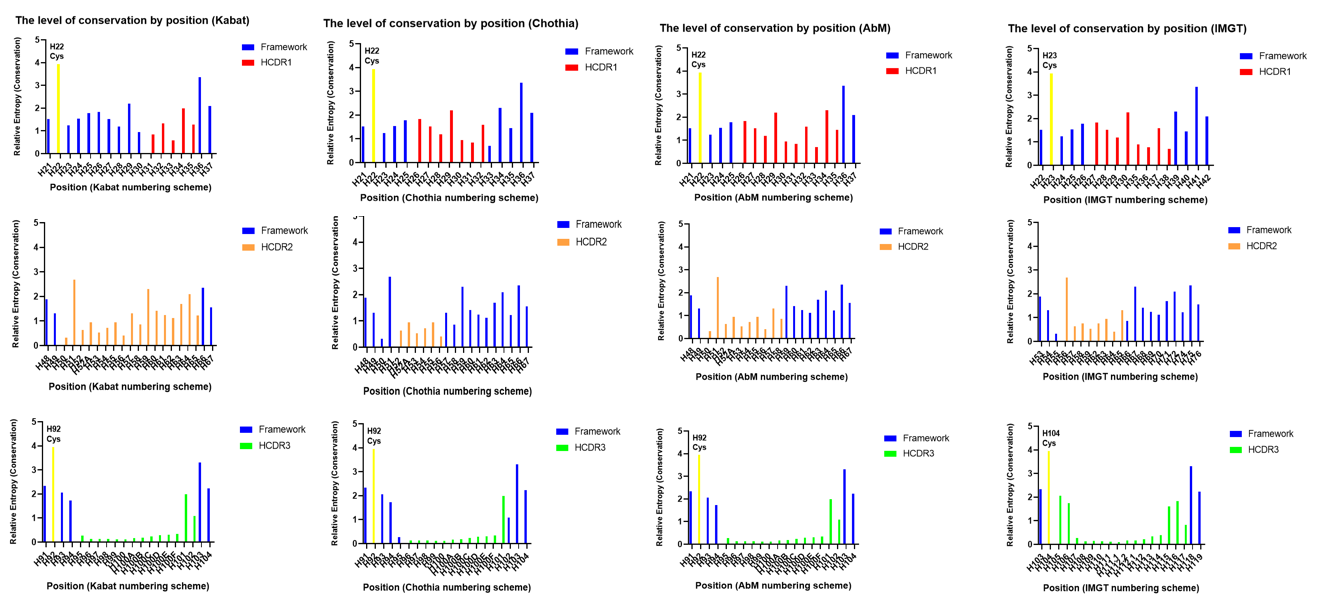

**Supplemental Figure 1b. Relative entropies of conservation for selected residues in the variable heavy domain.** The relative entropy of conservation was calculated for each residue, with residue position given by the Kabat, Chothia, AbM and IMGT numbering schemes. Relative entropies of residues within the CDR are colored according to the HCDR to which they belong (HCDR1 shown in red, HCDR2 in orange, and HCDR3 in green), while framework residues are shown in blue. Highlighted cysteine residues are shown in yellow.

**Figure S2a**

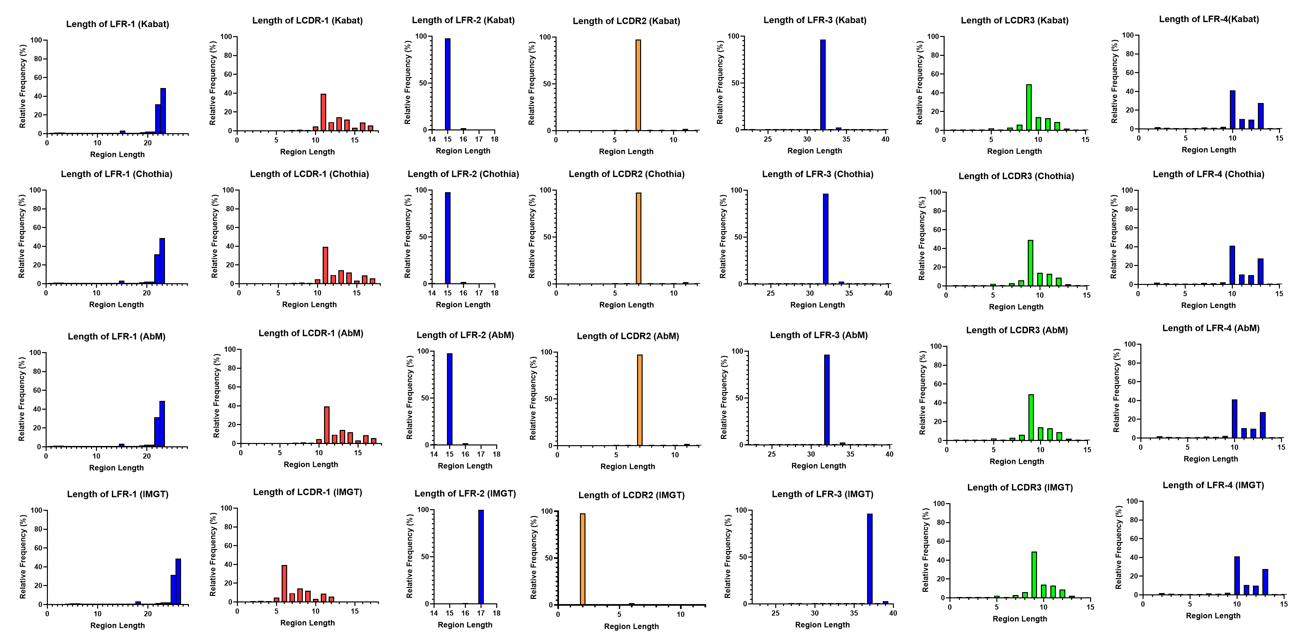

**Supplemental Figure 2a. Distribution of LCDR length for numbering schemes.** The length distribution of the CDR and framework regions in the variable light domain, as defined by the Kabat, Chothia, AbM and IMGT numbering schemes, are compared.

**Figure S2b**
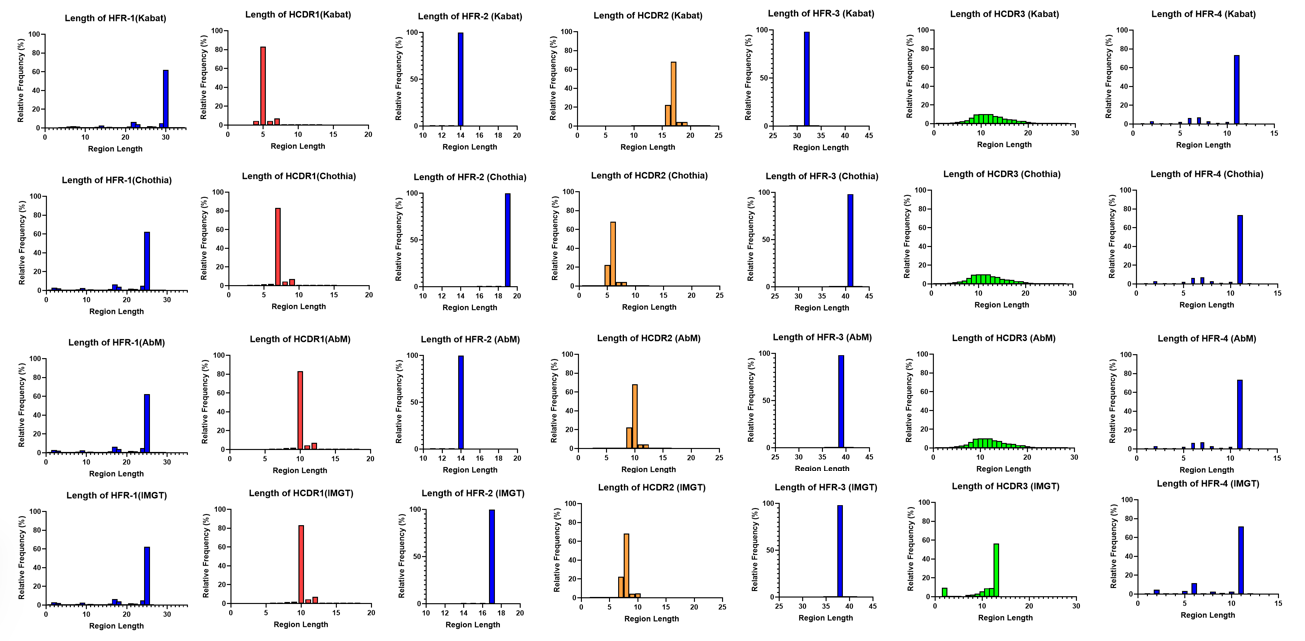

**Supplemental Figure 2b.** **Distribution of HCDRs length for numbering schemes.** The length distribution of the CDR and framework regions in the variable heavy domains, as defined by the Kabat, Chothia, AbM and IMGT numbering schemes, are compared.

**Figure S3**

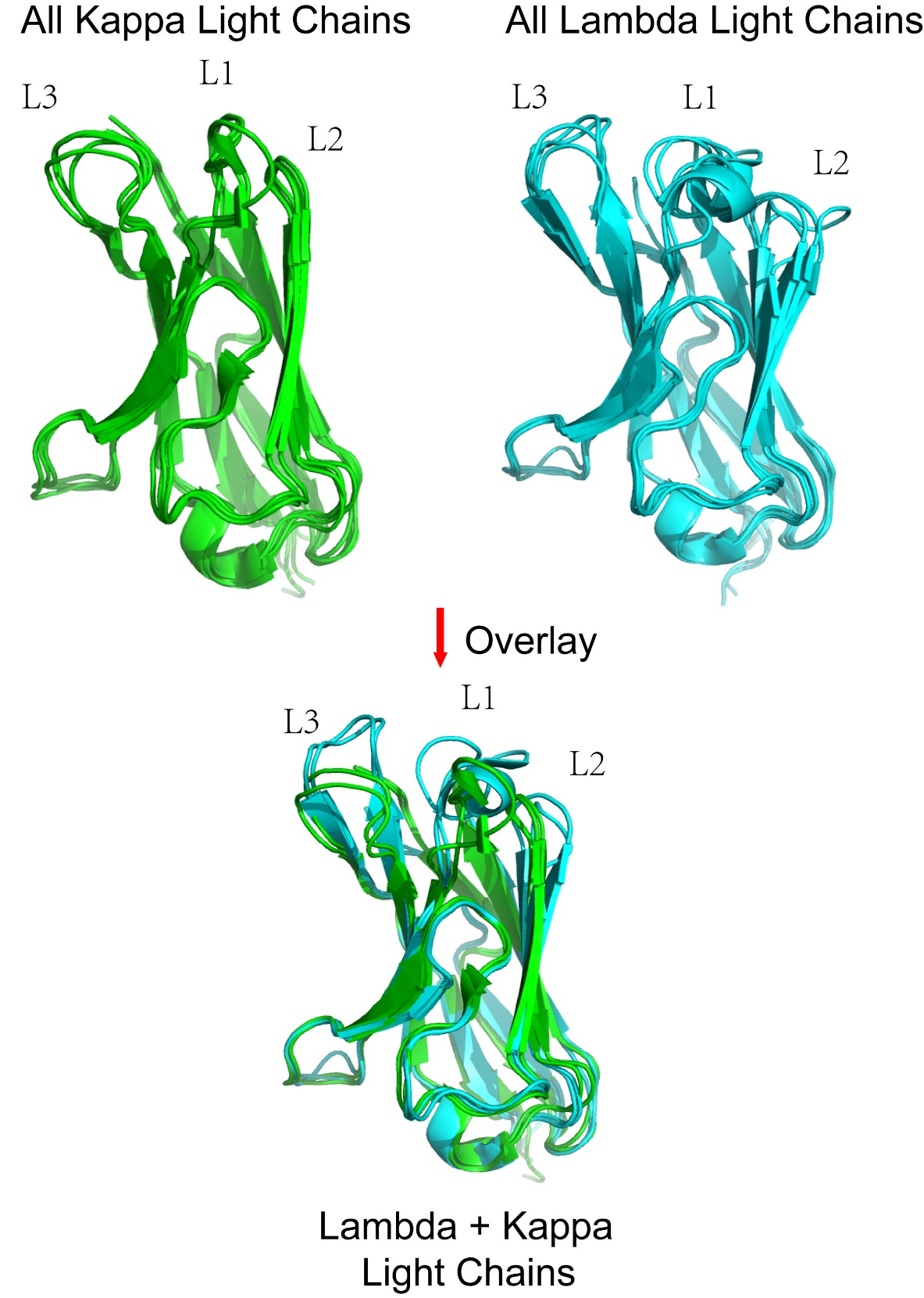

**Supplemental Figure 3. Structural comparison of the lambda and kappa light chains.** Shown is the structural alignment of the lambda light chain variable domains (Cyan, PDB ID: 6QBC, 5OD0, 4YNY, 6XRJ), the lambda light chain variable domains (Green, PDB ID: 2ZKH, 6TCM, 6Z7X, 5VH3), and the overlay alignment map of the two kappa light chain variable domain and two lambda light variable domain structures (PDB ID: 6QBC, 5OD0, 2ZKH, 6TCM). The lambda light chains exhibit a substantial degree of homology to each other, as do the kappa light chains. However, notably lower levels of homology are observed between the lambda and kappa light chains, particularly within CDR loop regions.

**Figure S4**

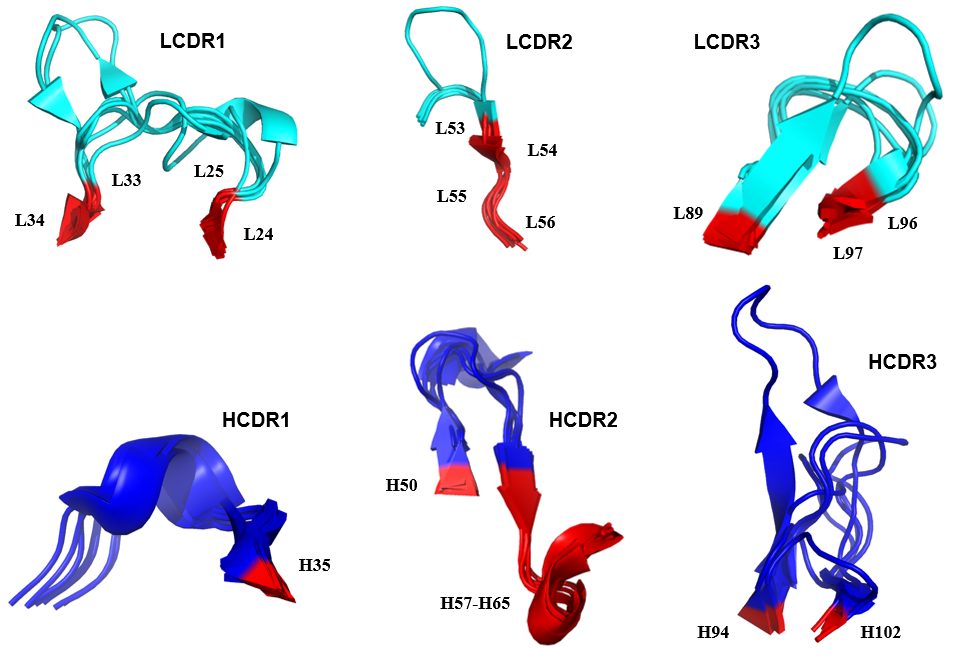

**Supplemental Figure 4. Structural alignment of the CDR loops from multiple antibody variable domains.** The structural alignment of the light chain variable domains and the heavy chain variable domains (PDB ID: 2ZKH, 6TCM, 6BE2, 6QBC, 6Z7X, 5VH3, 1CR9) are shown. CDR loops were defined according to the "general CDRs", which include all residues defined as CDRs by any numbering scheme and numbered according to the Kabat numbering scheme. Structurally conserved residues are color coded in red and their residue numbers annotated.

**Figure S5a**

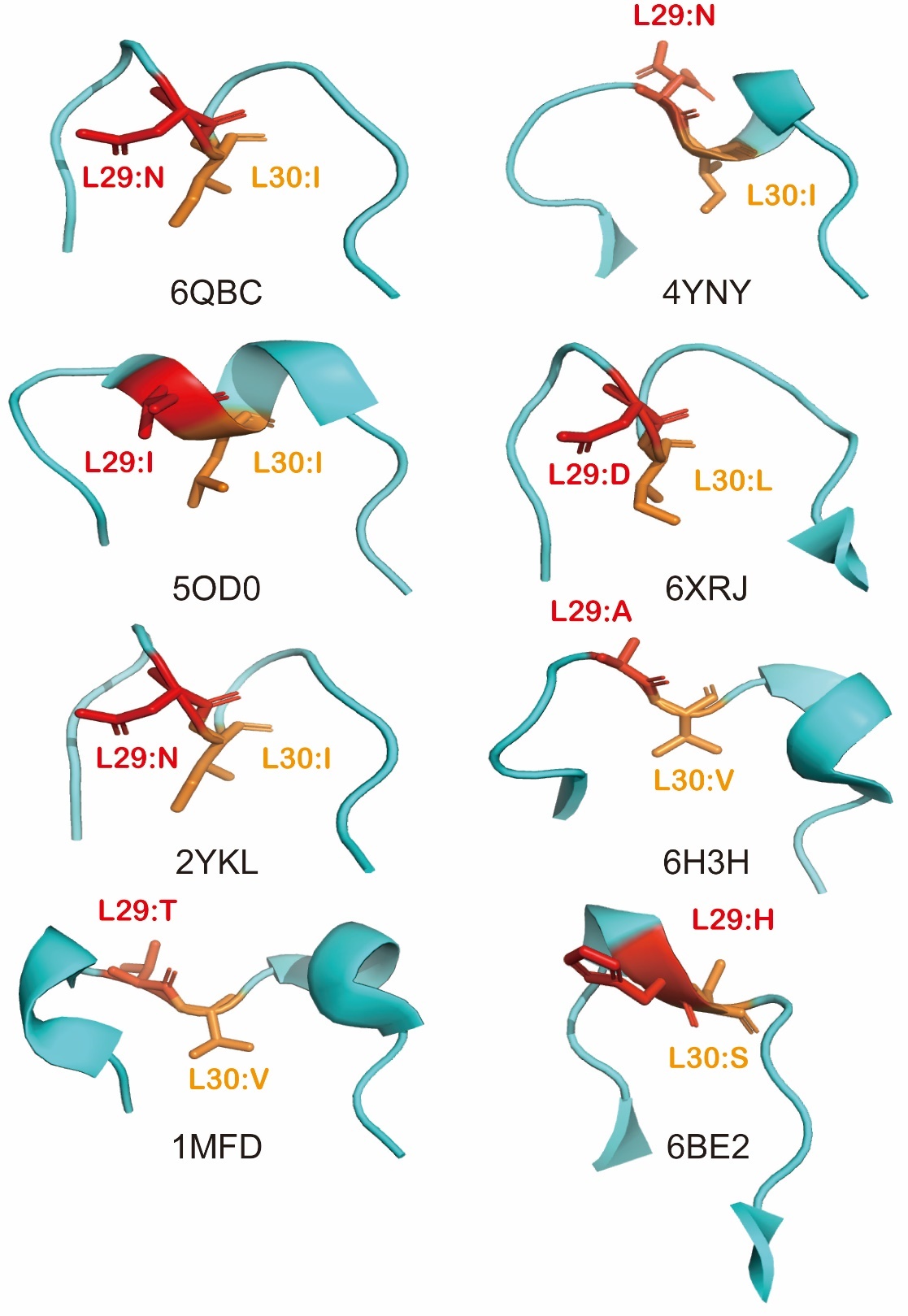

**Supplemental Figure 5a. A pivot point in the LCDR1 is located at L30 in the lambda light chain.** The LCDR1 loop structure of eight lambda light chain variable domains are shown (PDB ID: 6QBC, 4YNY, 5OD0, 6XRJ, 2YKL, 6H3H, 1MFD, 6BE2). The L29 (red) and L30 (orange) residues (IMGT numbering scheme) are annotated. Note that 6BE2 structure does not show an obvious pivot point feature in the loop—this may be due to the fact that it was derived from an IgG 2 variable domain.

**Figure S5b**

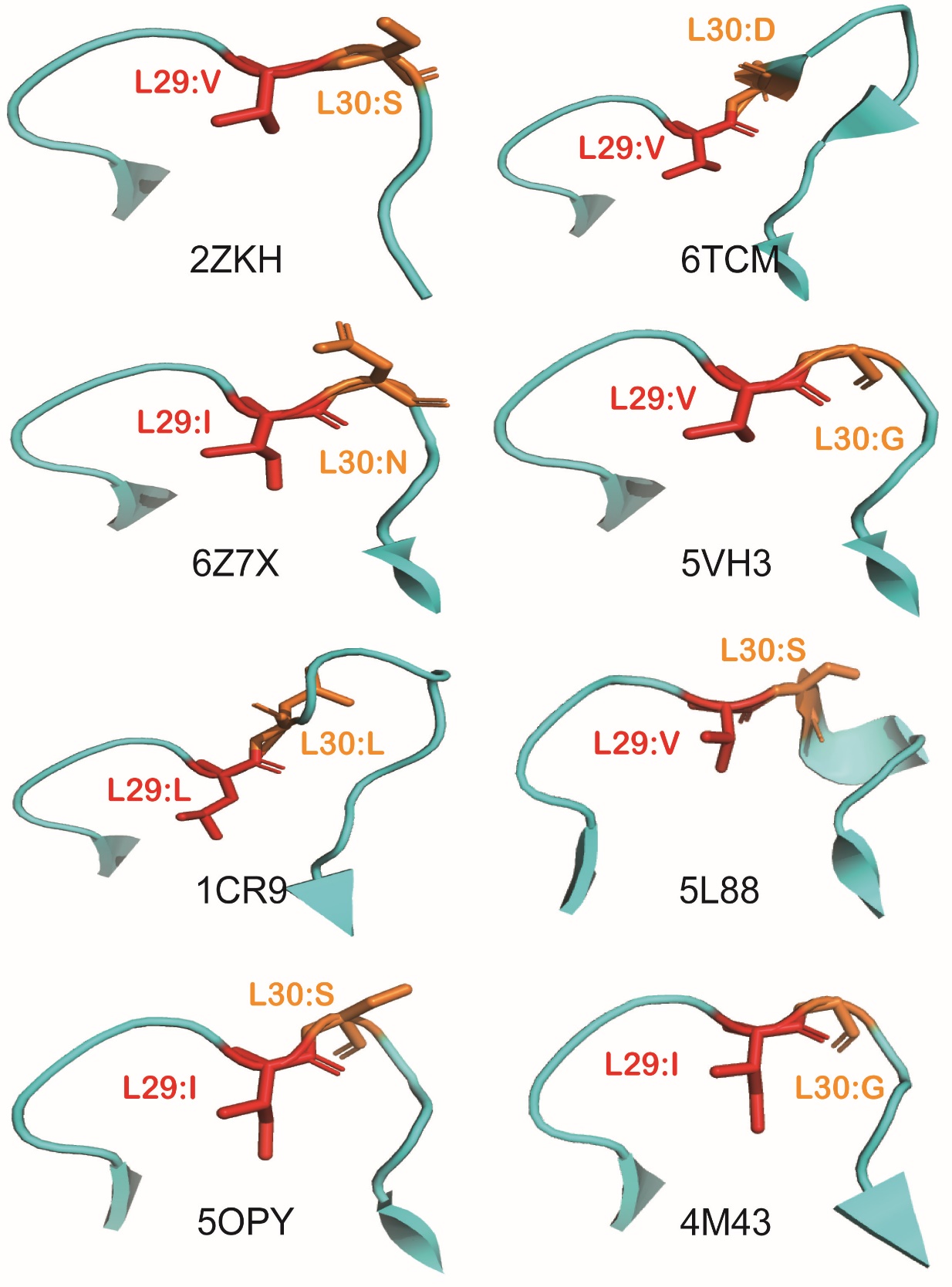

**Supplemental Figure 5b. Another pivot point in the LCDR1 is located at L29 in the kappa light chain.** The LCDR1 loop structure of eight kappa light chain variable domains are shown (PDB ID: 2ZKH, 6TCM, 6Z7X, 5VH3, 1CR9, 5L88, 5OPY, 4M43). The L29 (red) and L30 (orange) residues (IMGT numbering scheme) are annotated.

**Figure S6**

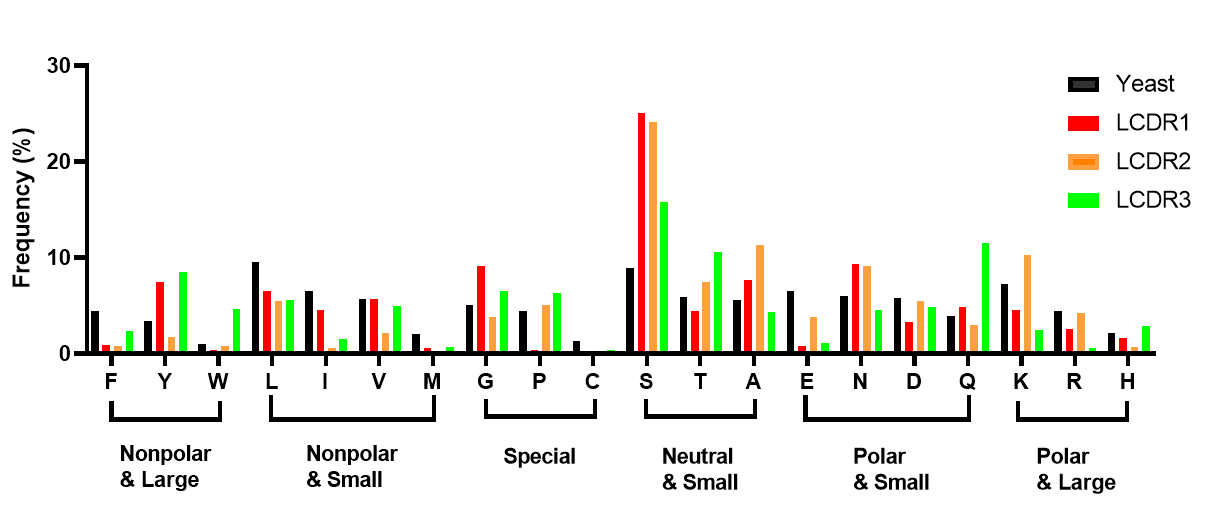

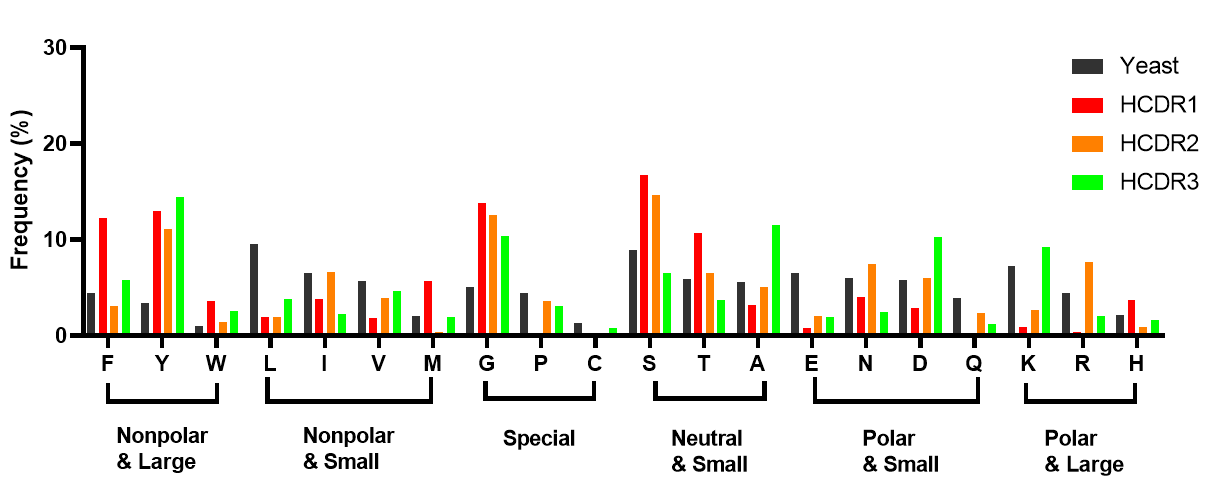

**Supplemental Figure 6. Amino acid distribution in each CDR loop compared to reference distribution in yeast.** CDR loops were defined according to the "general CDRs", which include all residues defined as CDRs by any numbering scheme (Kabat, Chothia, AbM and IMGT). The reference distribution “yeast” was generated based on codon usage in the yeast proteome.

**Figure S7**

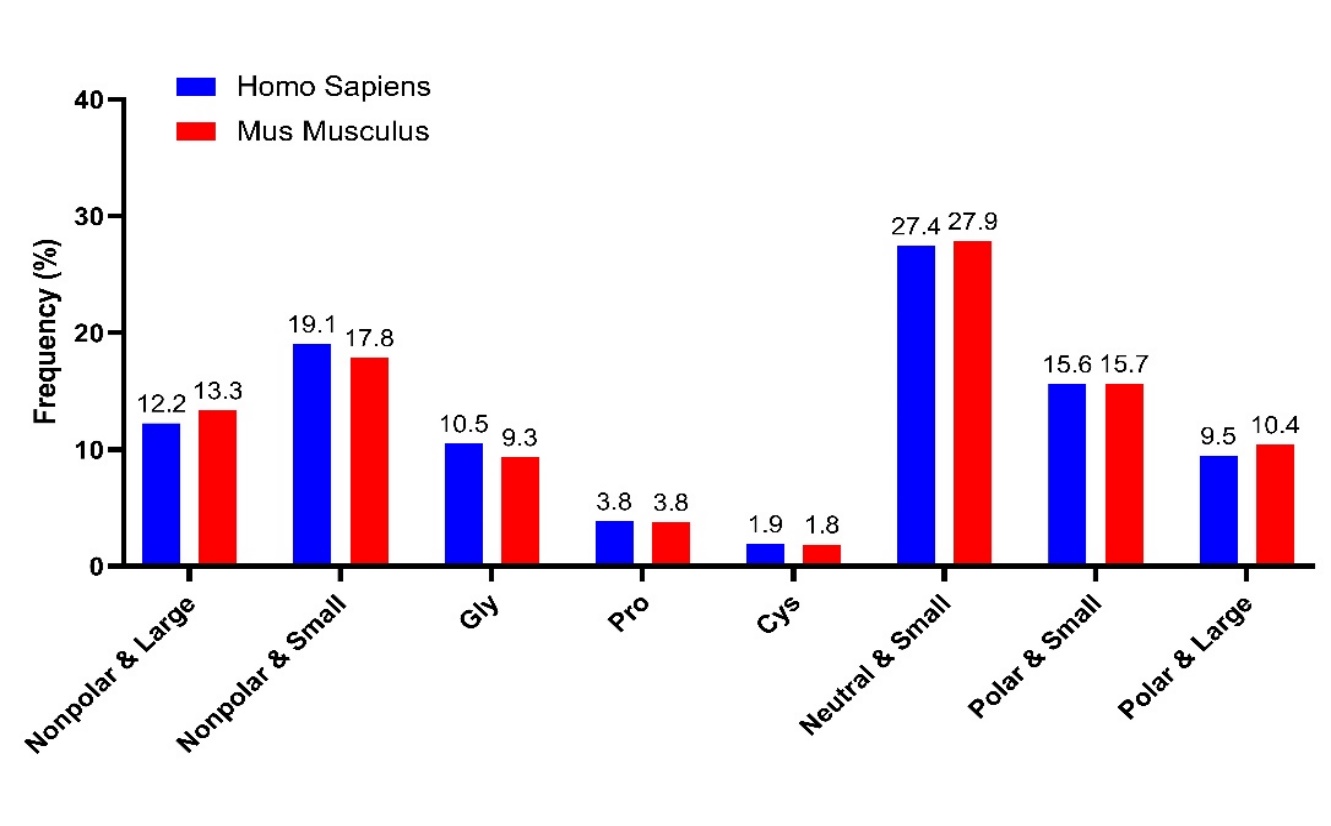

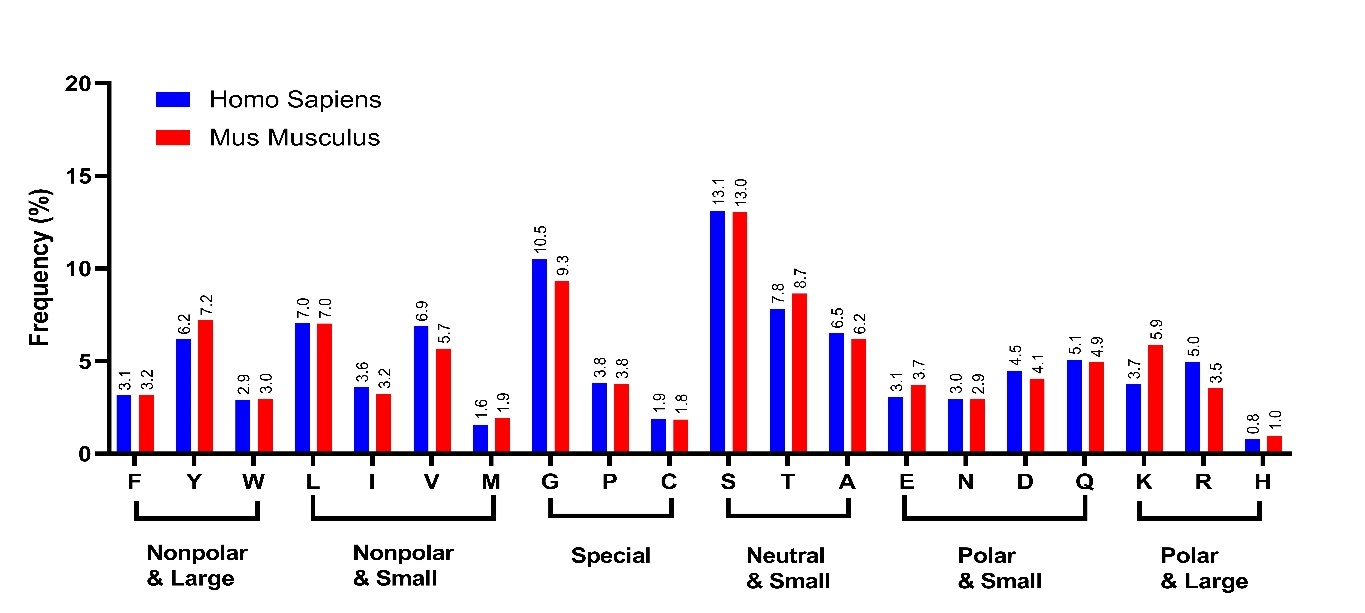

**Supplemental Figure 7. The amino acid distribution is similar for both mouse and human antibodies.** The amino acid distribution in the mouse and human antibody variable domains are compared, with amino acids grouped based on side chain volume and polarity.

**Figure S8**

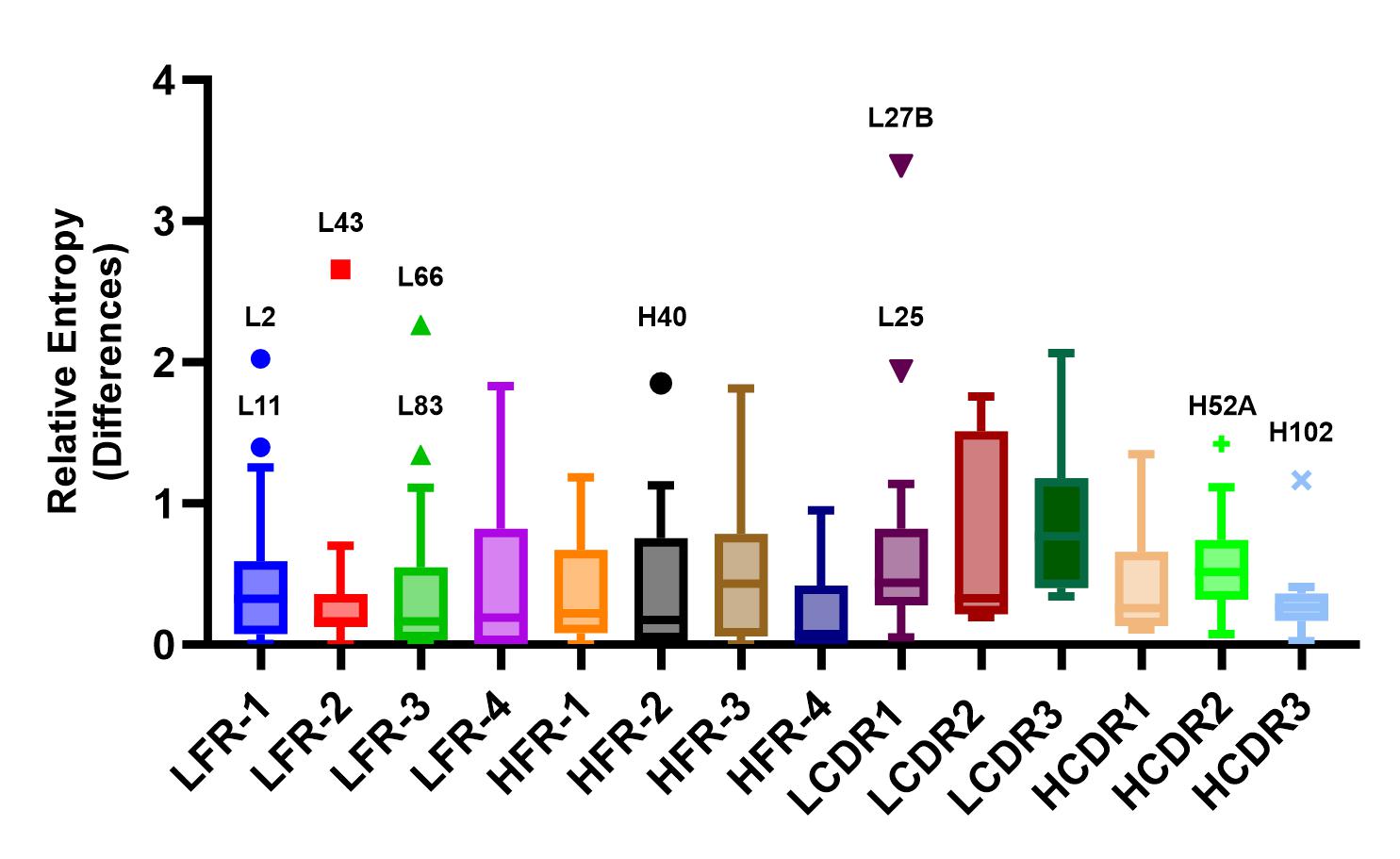

**Supplemental Figure 8. Box-and-whisker plot of the regional relative entropy differences between mouse and human variable domains.** CDRs are defined based on the inclusive “general CDR” definition and numbered in Kabat numbering scheme. Tukey boxplot was used to generate the plot, which calculates the inter-quartile distance (IQS, the difference between the 25th and 75th percentiles). The value between the 25^th^ and 75^th^ have been drawn as the box. The median is shown as a line within the box. Any data point falling beyond the 75th percentile plus 1.5 times the IQS was represented as an individual point on the plot, and the corresponding residue numbers were annotated above these points for reference.

**Figure S9**

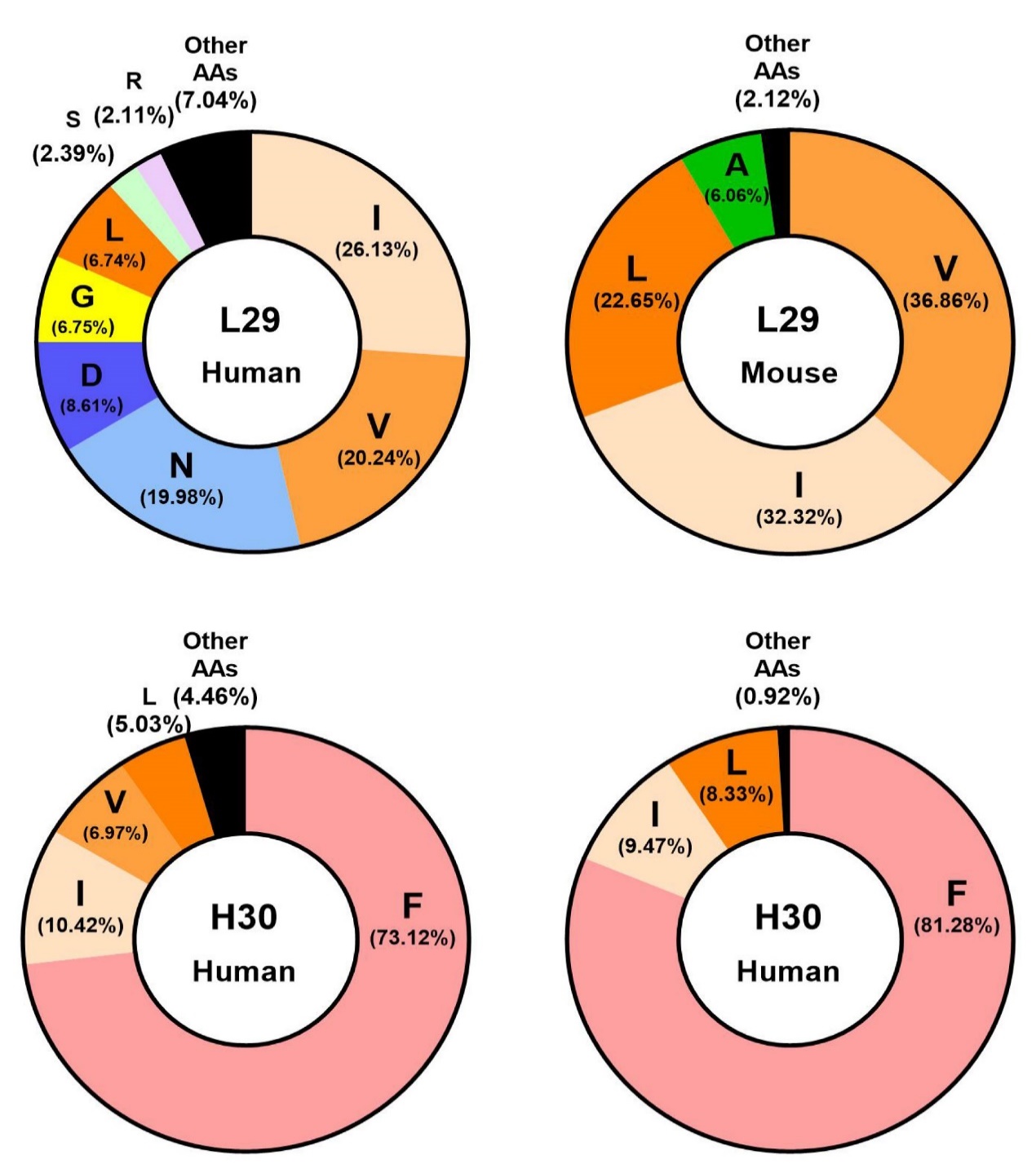

**Supplemental Figure 9. Amino acids distribution at pivot points in mouse and human antibody variable domains.** The amino acid distributions at the pivot points residues L29 and H30 (IMGT numbering scheme) are shown. Only amino acids exceeding a 2% occupancy rate are explicitly marked, while the frequencies of all other amino acids have been compiled and collectively represented as "Other AAs" in the figures for clarity.

**Figure S10**

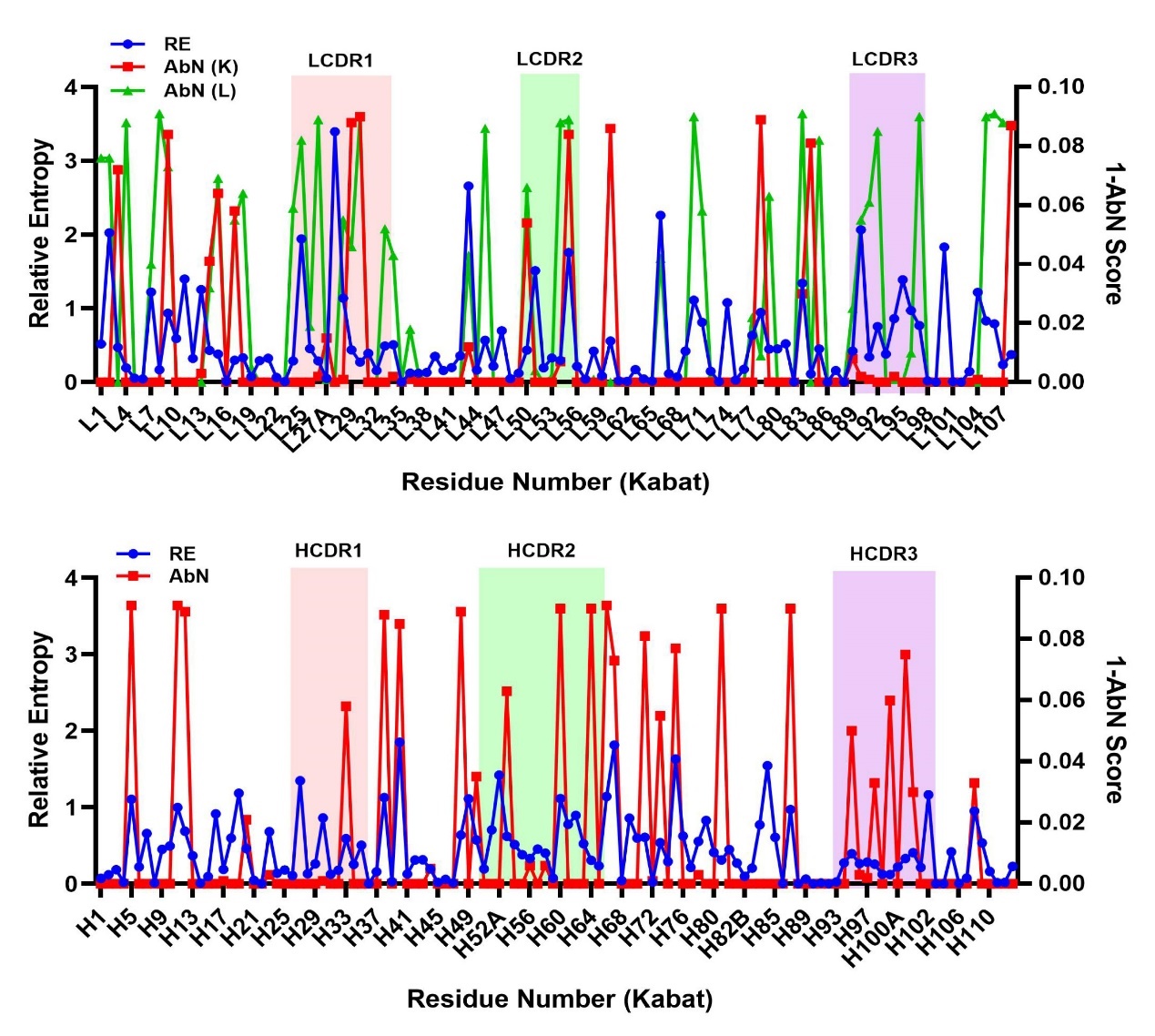

**Supplemental Figure 10.** **The residue wise Relative Entropy and the AbNatiV Score calculated based on the consensus Mus sequence in the variable domain** The value of RE and (1-AbNatiV score) of each residue in the variable light (top) and variable heavy (bottom) are shown in the graph. The relative entropy of differences between mouse and human species were shown in blue dots. The AbNatiV score of humanness were shown in red dots or green dots. In the variable light, the AvNatiV score were calculated based on the Variable light Kappa (red dots) and Variable light lambda (green dots) models. To align with the Relative entropy, the AbNatiV score were shown in (1-AbNatiV), which depict the non-humanness of the consensus Mus sequence in each of the position.

**Figure S11a**

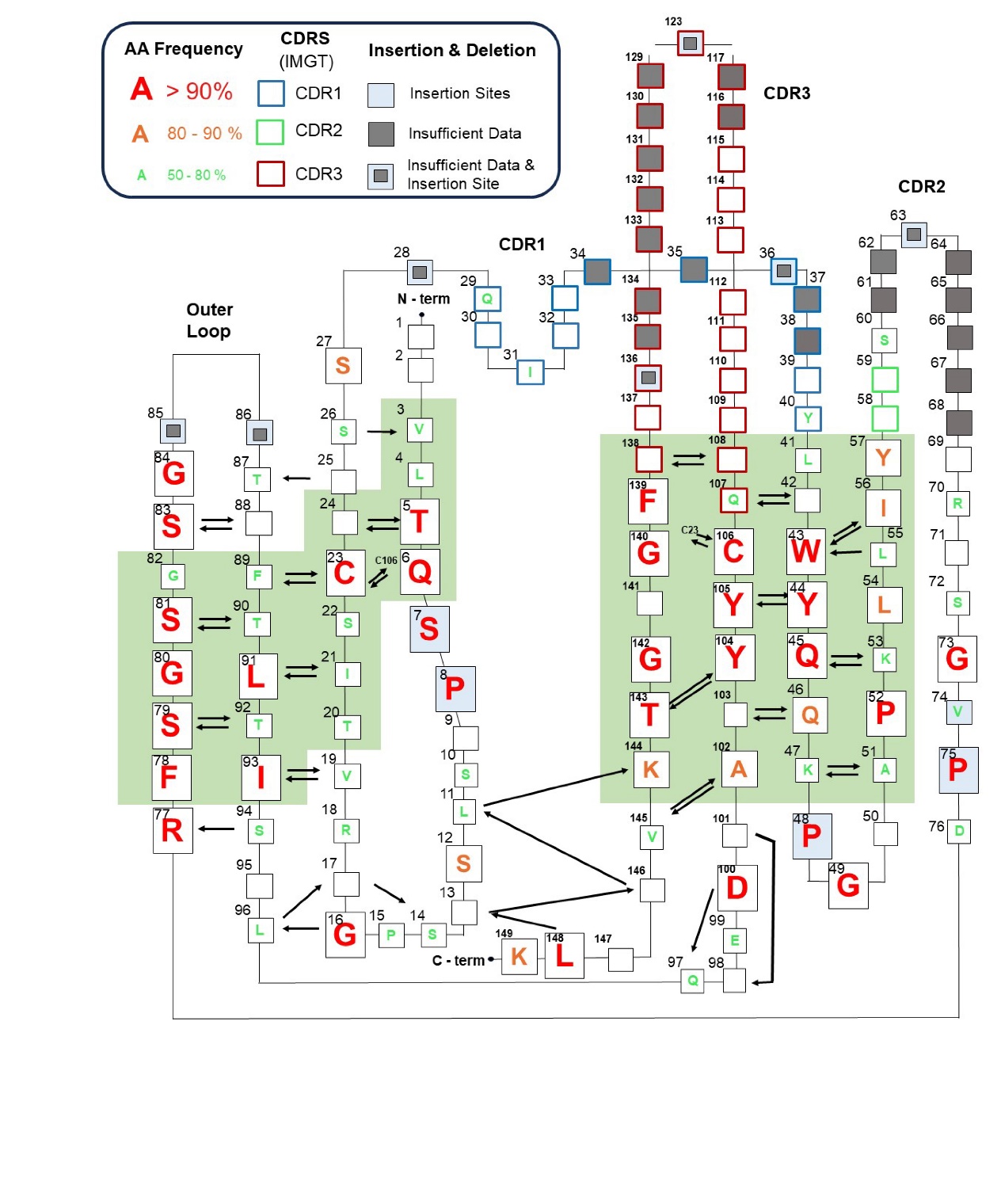

**Supplemental Figure 11a. A representation of the consensus sequence, consensus structure, and main-chain hydrogen bonding pattern of human immunoglobulin variable light domains.** A 2D map of the variable domain was generated based on the Honnegger numbering scheme. The residues were numbered based on the Aho numbering schemes. Complementarity Determining Regions (CDRs) are defined as per the IMGT numbering scheme. In the map, one-letter codes highlighted in red, orange, and green correspond to amino acid frequencies above 90%, within the range of 80-90%, and between 50% and 80% respectively. CDR residues are marked based on border color, with CDR1 in blue, CDR2 in green, and CDR3 in red. Arrows indicate hydrogen bonds present in the majority of structures for all types of immunoglobulin variable domains. The loop and turn regions which accommodate gaps are indicated in gray. Green areas underlie the residues with Cɑ positions that are highly structurally conserved.

**Figure S11b**

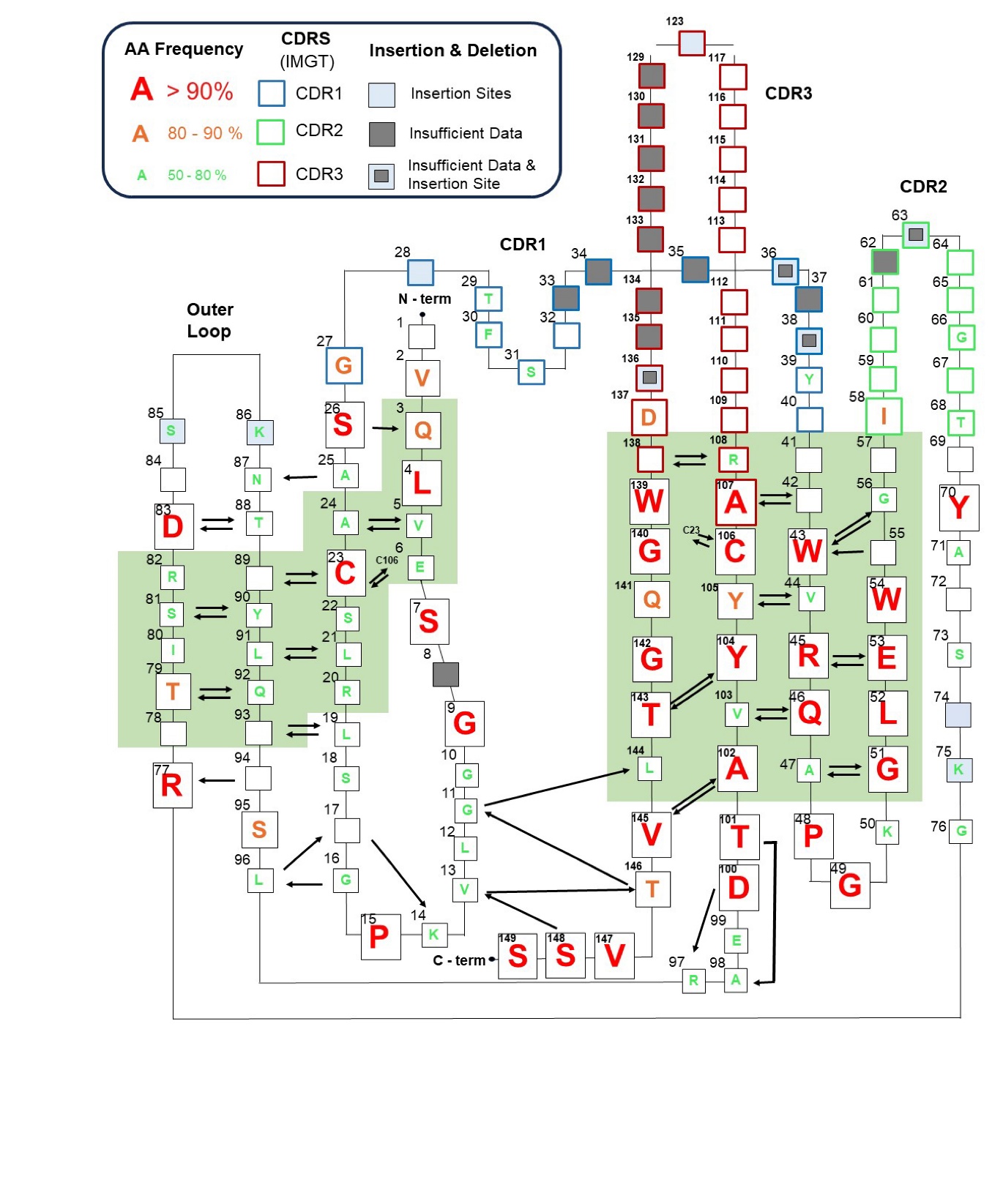

**Supplemental Figure 11b. A representation of the consensus sequence, consensus structure, and main-chain hydrogen bonding pattern of human immunoglobulin variable heavy domains.** A 2D map of the variable domain was generated based on the Honnegger numbering scheme. The residues were numbered in Aho numbering schemes. Complementarity Determining Regions (CDRs) are defined as per the IMGT numbering scheme. In the map, one-letter codes highlighted in red, orange, and green correspond to amino acid frequencies above 90%, within the range of 80-90%, and between 50% and 80% respectively. CDR residues are marked based on border color, with CDR1 in blue, CDR2 in green, and CDR3 in red. Arrows indicate hydrogen bonds present in the majority of the structures for all types of immunoglobulin variable domains. The loop and turn regions which accommodate gaps are indicated in gray. Green areas underlie the residues with Cɑ positions that are highly structurally conserved.

**SUPPLIEMENTARY TABLE 1**

Table of Relative Entopy Calculated in each Regions in the Variable Domains

| **Region** | **Avg. RE** (Conservation) | **Region** | **Avg. RE** (Conservation) |
| --- | --- | --- | --- |
| LFR-1 | 1.648 | HFR-1 | 1.801 |
| LFR-2 | 2.177 | HFR-2 | 2.326 |
| LFR-3 | 2.013 | HFR-3 | 1.891 |
| LFR-4 | 1.906 | HFR-4 | 2.118 |
| LCDR1 | 1.035 | HCDR1 | 1.205 |
| LCDR2 | 1.131 | HCDR2 | 1.204 |
| LCDR3 | 0.758 | HCDR3 | 0.394 |
| L Framework | 1.943 | H Framework | 1.959 |
| L CDRs | 0.923 | H CDRs | 0.889 |
| Variable Light Domain | 1.656 | Variable Heavy Domain | 1.646 |
| Variable Domains  Total | 1.651 | | |

**Supplemental Table 1.** The average Relative Entropy of Conservation calculated in each region, with CDRs annotated based on the Kabat Numbering Scheme.

**SUPPLIEMENTARY TABLE 2**

Comparison of the CDR definition of different numbering schemes for immunoglobulin variable domain

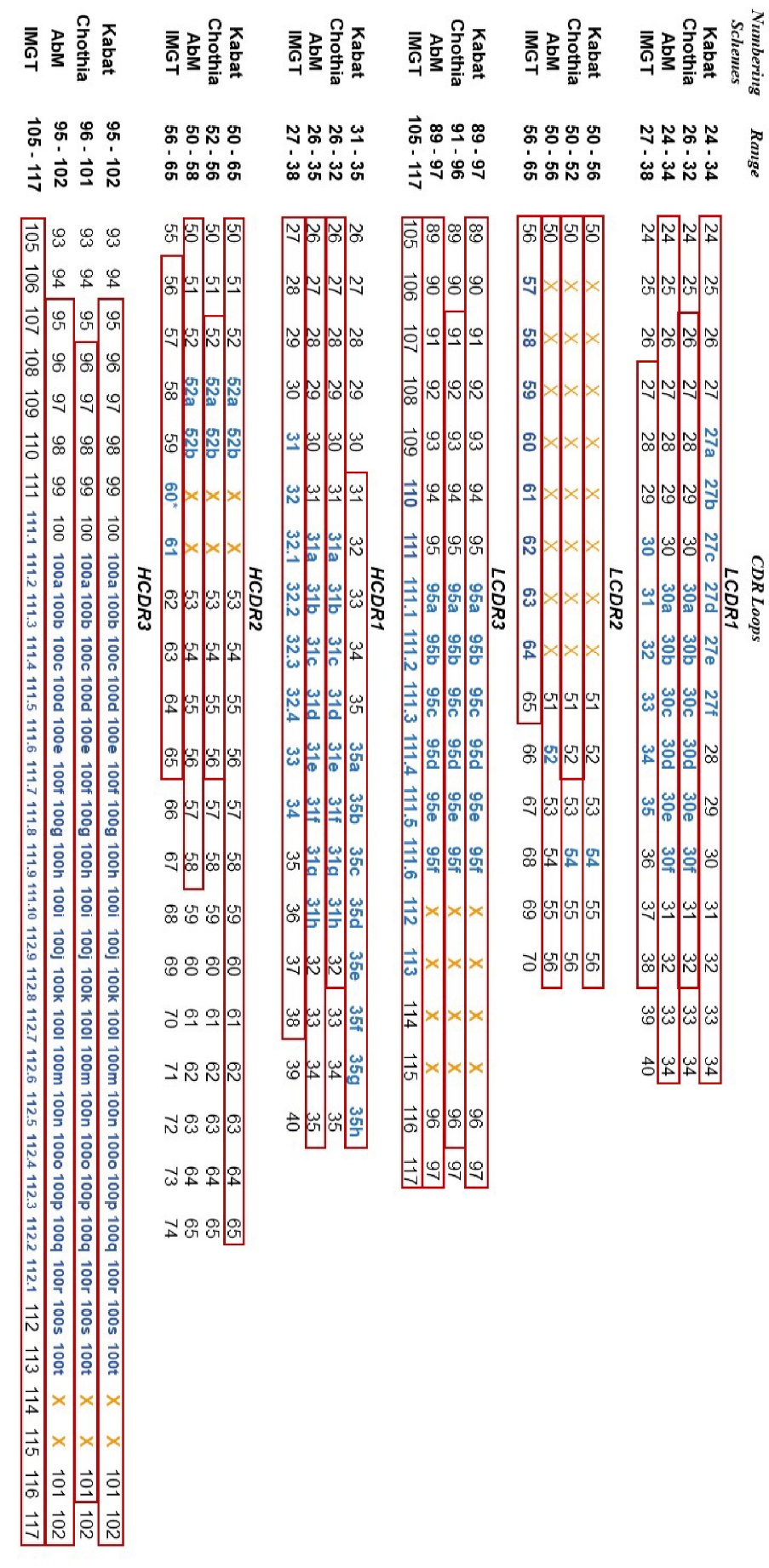

**Supplemental Table 2.** The different numbering schemes for immunoglobulin variable domains (Kabat, Chothia, AbM and IMGT) have been aligned (15, 17, 20, 21, 23, 34). CDR residues were defined based on previously published literature and highlighted within red frames (15, 17, 19, 20, 23). It's worth noting that "Chothia" in this context refers to the Consensus Chothia CDR definition developed by the Martin group (19). Inserted residues have been denoted in **bold** and appear in **blue** text. Residues that lack equivalent counterparts in the respective numbering scheme are indicated by an orange cross “**X**” in the table.

**SUPPLIEMENTARY TABLE 3a**

Table of Antibody Template PDB Entries, Organism, Light Chain Type and Reference Information

| **PDB ID** | **Organism** | **Light Chain Type** | **Reference** |
| --- | --- | --- | --- |
| ***Structure Alignment Analysis & Pivot Point Sequential Analysis*** | | | |
| 2ZKH | Homo Sapiens | **Kappa** |  |
| 6TCM | Homo Sapiens | **Kappa** |  |
| 6BE2 | Homo Sapiens | ***Lambda*** |  |
| 6QBC | Homo Sapiens | ***Lambda*** |  |
| 6Z7X | Mus Musculus | **Kappa** |  |
| 5VH3 | Mus Musculus | **Kappa** |  |
| 1CR9 | Mus Musculus | **Kappa** |  |
| ***Pivot Point Sequential Analysis*** | | | |
| 4YNY | Rattus norvegicus | ***Lambda*** |  |
| 5OD0 | Homo Sapiens | ***Lambda*** |  |
| 6XRJ | Homo Sapiens | ***Lambda*** |  |
| 2YKL | Homo Sapiens | ***Lambda*** |  |
| 6H3H | Mus Musculus | ***Lambda*** |  |
| 1MFD | Mus Musculus | ***Lambda*** |  |
| 5L88 | Mus Musculus | **Kappa** |  |
| 5OPY | Mus Musculus | **Kappa** |  |
| 4M43 | Mus Musculus | **Kappa** |  |

**Supplemental Table 3a.** The PDB ID, organism and light chain type information of the immunoglobulin used for structural alignment.

**SUPPLIEMENTARY TABLE 3b**

Table of Antibody Template Protein Amino Acid Sequences

| **PDB ID** | **Sequence** |
| --- | --- |
| 2ZKH | **Light Chain**  QVVLTQSPGIMSASPGEKVTITCSASSSVSYMYWFQQKPGTSPKLWIYSTSNLASGVPARFRGSGSGTSYSLTISRMEAEDAATYYCQQRSGYPRTFGGGTKLEIKRA  *DAAPTVSIFPPSSEQLTSGGASVVCFLNNFYPKDINVKWKIDGSERQNGVLNSWTDQDSKDSTYSMSSTLTLTKDEYERHNSYTCEATHKTSTSPIVKSFNRNEC*  **Heavy Chain**  EVKLEESGGGLVQPGGSMKLSCAASGFTFSDAWMDWVRQSPEKGLEWVAEIRSKVNNHAIHYAESVKGRFTVSRDDSKSSVYLQMNSLRAEDTGIYYCSGWSFLYWGQGTLVTVSA  *AKTTPPSVYPLAPGSAAQTNSMVTLGCLVKGYFPEPVTVTWNSGSLSSGVHTFPAVLQSDLYTLSSSVTVPSSTWPSETVTCNVAHPASSTKVDKKIVPRD* |
| 6TCM | **Light Chain**  DIQLTQSPSSLSASVGDRVTITCRASQSVDYDGDSYMNWYQQKPGKAPKLLIYAASYLESGVPSRFSGSGSGTDFTLTISSLQPEDFATYYCQQSHEDPYTFGQGTKVEIKRTV  *AAPSVFIFPPSDEQLKSGTASVVCLLNNFYPREAKVQWKVDNALQSGNSQESVTEQDSKDSTYSLSSTLTLSKADYEKHKVYACEVTHQGLSSPVTKSFNRGEC*  **Heavy Chain**  EVQLVESGGGLVQPGGSLRLSCAVSGYSITSGYSWNWIRQAPGKGLEWVASITYDGSTNYNPSVKGRITISRDDSKNTFYLQMNSLRAEDTAVYYCARGSHYFGHWHFAVWGQGTLVTVSS  *ASTKGPSVFPLAPSSKSTSGGTAALGCLVKDYFPEPVTVSWNSGALTSGVHTFPAVLQSSGLYSLSSVVTVPSSSLGTQTYICNVNHKPSNTKVDKKVEPKSCHHHHHH* |
| 6BE2 | **Light Chain**  QLVLTQSPSASASLGASVKLTCTLSSGHSNYAIAWHQQQPGKGPRYLMKVNRDGSHIRGDGIPDRFSGSTSGAERYLTISSLQSEDEADYYCQTWGAGIRVFGGGTKLTVLGQP  *KAAPSVTLFPPSSEELQANKATLVCLISDFYPGAVTVAWKADGSPVKAGVETTTPSKQSNNKYAASSYLSLTPEQWKSHRSYSCQVTHEGSTVEKTVAPTECS*  **Heavy Chain**  QVQLQESGPGLVKPSETLSLTCTVSGGSISGYYWSWIRQPPGKGLEWIGYIHYSRSTNSNPALKSRVTISSDTSKNQLSLRLSSVTAADTAVYYCARDTYYYDSGDYEDAFDIWGQGTMVTVSS  *ASTKGPSVFPLAPSSKSTSGGTAALGCLVKDYFPEPVTVSWNSGALTSGVHTFPAVLQSSGLYSLSSVVTVPSSSLGTQTYICNVNHKPSNTKVDKKAEPKSC* |
| 6QBC | **Light Chain**  QSVLTQPPSASGTPGQRVTISCSGSSSNIGSNTVNWYQQLPGTAPKLLIYSNNQRPSGVPDRFSGSKSGTSASLAISGLQSEDEADYYCAAWDDSLNAWVFGGGTKLTVLGES  *EGQPKSSPSVTLFPPSSEELETNKATLVCTITDFYPGVVTVDWKVDGTPVTQGMETTQPSKQSNNKYMASSYLTLTARAWERHSSYSCQVTHEGHTVEKSLSRADCS*  **Heavy Chain**  QVTLKESGGGLVKPGGSLRLSCAASGFTFSSYSMNWVRQAPGKGLEWVSSISSSSSYIYYADSVKGRFTISRDNAKNSLYLQMNSLRAEDTAVYYCARQVGATWAFDIWGQGTLVTVSA  *AKTTPPSVYPLAPGSAAQTNSMVTLGCLVKGYFPEPVTVTWNSGSLSSGVHTFPAVLQSDLYTLSSSVTVPSSPRPSETVTCNVAHPASSTKVDKKIVPRDCAAAENLYFQ* |
| 6Z7X | **Light Chain**  DIQMTQSPSSLSASLGGRVTITCKASQDINKYLAWYQHKPGKGPRLLIHYTSTLQPGIPSRFSGSGSGRDYSFSISNLEPEDVATYYCLQYDSLLSFGAGTKLELKRA  *DAAPTVSIFPPSSEQLTSGGASVVCFLNNFYPKDINVKWKIDGSERQNGVLNSWTDQDSKDSTYSMSSTLTLTKDEYERHNSYTCEATHKTSTSPIVKSFNRNEC*  **Heavy Chain**  EVQLVESGGGLVKPGGSLKLSCTASGFAFSDYDMSWVRQTPEKRLEWVAFISNGGYSTYYPDTVKGRFTISRDNAENTLYLQMSSLKSEDTAIYYCARQGLRYFDYWGLGTTLTVSS  *AKTTPPSVYPLAPGSAAQTNSMVTLGCLVKGYFPEPVTVTWNSGSLSSGVHTFPAVLQSDLYTLSSSVTVPSSTWPSETVTCNVAHPASSTKVDKKIVPRDCG* |
| 5VH3 | **Light Chain**  DILLTQSPAILSVSPGERVSFSCRASQFVGSSIHWYQQRTNGSPRLLIKYASESMSGIPSRFSGSGSGTDFTLSINTVESEDIADYYCQQSHSWPFTFGSGTNLEVKRTV  *AAPSVFIFPPSDEQLKSGTASVVCLLNNFYPREAKVQWKVDNALQSGNSQESVTEQDSKDSTYSLSSTLTLSKADYEKHKVYACEVTHQGLSSPVTKSFNRGEC*  **Heavy Chain**  EVKLEESGGGLVQPGGSMKLSCVASGFIFSNHWMNWVRQSPEKGLEWVAEIRSKSINSATHYAESVKGRFTISRDDSKSAVYLQMTDLRTEDTGVYYCSRNYYGSTYDYWGQGTTLTVSS  *ASTKGPSVFPLAPSSKSTSGGTAALGCLVKDYFPEPVTVSWNSGALTSGVHTFPAVLQSSGLYSLSSVVTVPSSSLGTQTYICNVNHKPSNTKVDKKVEPKSCDKT* |
| 1CR9 | **Light Chain**  DVVMTQTPLSLSVTIGQPASISCKSSQSLLDSDGKTYLIWVFQRPGQSPKRLIFLVSKRDSGVPDRFTGSGSGTDFTLKISRVEAEDVGVYYCWQGTHFPHTVGGGTKLEIARAD  *AAPTVSIFPPSSEQLTSGGASVVCFLNNFYPKDINVKWKIDGSERQNGVLNSWTDQDSKDSTYSMSSTLTLTKDEYERHNSYTCEATHKTSTSPIVKSFNRNEC*  **Heavy Chain**  KVKLQQSGAELVRSGASVKLSCTASGFNIKDYYIQWVKQRPEQGLEWIGWIDPENGNSEYAPRFQGKATMTADTLSNTAYLQLSSLTSEDTAVYYCNADLHDYWGQGTTLTVSS  *AKTTAPSVYPLAPVCGDTTGSSVTLGCLVKGYFPEPVTLTWNSGSLSSGVHTFPAVLQSDLYTLSSSVTVTSSTWPSQSITCNVAHPASSTKVDKKIEPRVTS* |

**Supplemental Table 3b.** The amino acid sequences of the immunoglobulin used in the structure alignment. Sequences belonging to constant domains are *italicized*.

| PDB ID | Numbering schemes | LCDR1 | LCDR2 | LCDR3 |
| --- | --- | --- | --- | --- |
| 2ZKH | Kabat | SASSSVSYMY | STSNLAS | QQRSGYPRT |
|  | Chothia | SSSVSY | STS | RSGYPR |
|  | AbM | SASSSVSYMY | STSNLAS | QQRSGYPRT |
|  | IMGT | SSVSY | ST | QQRSGYPRT |
| 6TCM | Kabat | RASQSVDYDGDSYMN | AASYLES | QQSHEDPYT |
|  | Chothia | SQSVDYDGDSY | AAS | SHEDPY |
|  | AbM | RASQSVDYDGDSYMN | AASYLES | QQSHEDPYT |
|  | IMGT | QSVDYDGDSY | AA | QQSHEDPYT |
| 6BE2 | Kabat | TLSSGHSNYAIA | VNRDGSHIRGD | QTWGAGIRV |
|  | Chothia | SSGHSNYA | VNR | WGAGIR |
|  | AbM | TLSSGHSNYAIA | VNRDGSHIRGD | QTWGAGIRV |
|  | IMGT | SGHSNYA | VNRDGS | QTWGAGIRV |
| 6QBC | Kabat | SGSSSNIGSNTVN | SNNQRPS | AAWDDSLNAWV |
|  | Chothia | SSSNIGSNT | SNN | WDDSLNAW |
|  | AbM | SGSSSNIGSNTVN | SNNQRPS | AAWDDSLNAWV |
|  | IMGT | SSNIGSNT | SN | AAWDDSLNAWV |
| 6Z7X | Kabat | KASQDINKYLA | YTSTLQP | LQYDSLLS |
|  | Chothia | SQDINKY | YTS | YDSLL |
|  | AbM | KASQDINKYLA | YTSTLQP | LQYDSLLS |
|  | IMGT | QDINKY | YT | LQYDSLLS |
| 5VH3 | Kabat | RASQFVGSSIH | YASESMS | QQSHSWPFT |
|  | Chothia | SQFVGSS | YAS | SHSWPF |
|  | AbM | RASQFVGSSIH | YASESMS | QQSHSWPFT |
|  | IMGT | QFVGSS | YA | QQSHSWPFT |
| 1CR9 | Kabat | KSSQSLLDSDGKTYLI | LVSKRDS | WQGTHFPHT |
|  | Chothia | SQSLLDSDGKTY | LVS | GTHFPH |
|  | AbM | KSSQSLLDSDGKTYLI | LVSKRDS | WQGTHFPHT |
|  | IMGT | QSLLDSDGKTY | LV | WQGTHFPHT |

**SUPPLIEMENTARY TABLE 4a**

Table of Antibody Template Protein LCDRs Amino Acid Sequences

**Supplemental Table 4a.** The amino acid sequences of the LCDRs predicted by the Kabat, Chothia, AbM (Martin) and IMGT numbering schemes.

**SUPPLIEMENTARY TABLE 4b**

Table of Antibody Template Protein HCDRs Amino Acid Sequences

| PDB ID | Numbering schemes | HCDR1 | HCDR2 | HCDR3 |
| --- | --- | --- | --- | --- |
| 2ZKH | Kabat | DAWMD | EIRSKVNNHAIHYAESVKG | WSFLY |
|  | Chothia | GFTFSDA | RSKVNNHA | WSFLY |
|  | AbM | GFTFSDAWMD | EIRSKVNNHAIH | WSFLY |
|  | IMGT | GFTFSDAW | IRSKVNNHAI | SGWSFLY |
| 6TCM | Kabat | SGYSWN | SITYDGSTNYNPSVKG | GSHYFGHWHFAV |
|  | Chothia | GYSITSGY | TYDGS | GSHYFGHWHFAV |
|  | AbM | GYSITSGYSWN | SITYDGSTN | GSHYFGHWHFAV |
|  | IMGT | GYSITSGYS | ITYDGST | ARGSHYFGHWHFAV |
| 6BE2 | Kabat | GYYWS | YIHYSRSTNSNPALKS | DTYYYDSGDYEDAFDI |
|  | Chothia | GGSISGY | HYSRS | DTYYYDSGDYEDAFDI |
|  | AbM | GGSISGYYWS | YIHYSRSTN | DTYYYDSGDYEDAFDI |
|  | IMGT | GGSISGYY | IHYSRST | ARDTYYYDSGDYEDAFDI |
| 6QBC | Kabat | SYSMN | SISSSSSYIYYADSVKG | QVGATWAFDI |
|  | Chothia | GFTFSSY | SSSSSY | QVGATWAFDI |
|  | AbM | GFTFSSYSMN | SISSSSSYIY | QVGATWAFDI |
|  | IMGT | GFTFSSYS | ISSSSSYI | ARQVGATWAFDI |
| 6Z7X | Kabat | DYDMS | FISNGGYSTYYPDTVKG | QGLRYFDY |
|  | Chothia | GFAFSDY | SNGGYS | QGLRYFDY |
|  | AbM | GFAFSDYDMS | FISNGGYSTY | QGLRYFDY |
|  | IMGT | GFAFSDYD | ISNGGYST | ARQGLRYFDY |
| 5VH3 | Kabat | NHWMN | EIRSKSINSATHYAESVKG | NYYGSTYDY |
|  | Chothia | GFIFSNH | RSKSINSA | NYYGSTYDY |
|  | AbM | GFIFSNHWMN | EIRSKSINSATH | NYYGSTYDY |
|  | IMGT | GFIFSNHW | IRSKSINSAT | SRNYYGSTYDY |
| 1CR9 | Kabat | DYYIQ | WIDPENGNSEYAPRFQG | DLHDY |
|  | Chothia | GFNIKDY | DPENGN | DLHDY |
|  | AbM | GFNIKDYYIQ | WIDPENGNSE | DLHDY |
|  | IMGT | GFNIKDYY | IDPENGNS | NADLHDY |

**Supplemental Table 4b.** The amino acid sequences of the HCDRs predicted by the Kabat, Chothia, AbM (Martin) and IMGT numbering schemes.

**SUPPLIEMENTARY TABLE 5**

Table of Sequential and Structural Conserved Residues in CDRs

| **Loop** | **Kabat** | | **Chothia** | | **AbM** | | **IMGT** | |
| --- | --- | --- | --- | --- | --- | --- | --- | --- |
|  | **Seq.** | **Str.** | **Seq.** | **Str.** | **Seq.** | **Str.** | **Seq.** | **Str.** |
| LCDR1 | - | L24-L25  L32-L34 | - | - | - | L24-L25  L32-L34 | - | - |
| LCDR2 | L54 | L53-L56 | - | L50 | L54 | L53-L56 | - | 56 |
| LCDR3 | - | L89-L90  L97 | - | - | - | L89-L90  L97 | - | - |
| HCDR1 | - | H35 | - | - | - | H35 | 30 | - |
| HCDR2 | H51 | H50,  H57-H65 | - | - | H51 | H50,  H57-H58 | 56 | 65 |
| HCDR3 | H101 | H102 | H101 | - | H101 | H102 | 105,116 | 105, 117 |

**Supplemental Table 5.** The sequential (Seq.) and structural (Str.) conserved residues in CDRs identified based on each numbering scheme. To identify sequential conservation, residues with Relative Entropy (RE) values exceeding the average RE for the region by 1.65 times the standard deviation were flagged. Meanwhile, structural conservation was ascertained through structural alignment. Residues were annotated and numbered in accordance with each of the respective numbering schemes.

| **PDB ID** | **Sequence** |
| --- | --- |
| **Lambda Light Chains** | |
| 6QBC | QSVLTQPPSASGTPGQRVTISCSGSSS**NI**GSNTVNWYQQLPGTAPKLLIYSNNQRPSGVPDRFSGSKSGTSASLAISGLQSEDEADYYCAAWDDSLNAWVFGGGTKLTVLGES |
| 4YNY | QFVLTQPNSVSTNLGSTVKLSCKRSTG**NI**GSNYVNWYQQHEGRSPTTMIYRDDKRPDGVPDRFSGSIDRSSNSALLTINNVQTEDEADYFCHSYSSGIVFGGGTKLTVLGQP |
| 5OD0 | QSVWTQPPSVSAAPGQKVTISCSGDDS**I**LRSAFVSWYQQVPGSAPKLVIFDDRQRPSGIPARFSGSNSGTTATLDIAGLQRGDEADYYCAAWNGRLSAFVFGSGTKLTVLGQP |
| 6XRJ | ALTQPPSVSGSPGQSVIISCTGTSS**DI**GQYNSVSWYQQHPDKAPKLVIYGVTSRPSGVSDRFSGSKYGDTASLTISGLQAEDEADYYCSSHADENMALFGGGTRLTVLGQP |
| 2YKL | QSELTQPPSASGTPGQRVTISCSGSSS**NI**GSNYVYWYQQLPGTAPKLLIYRNNQRPSGVPDRFSGSKSGTSASLAISGLRSEDEADYYCAAWDDSLSAWVFGGGTQLDILGQP |
| 6H3H | QAVVTQESALTTSPGETVTLTCRSSTG**AV**TTSNYANWVQEKPDHLFTGLIGGTNNRAPGVPARFSGSLIGDKAALTITGAQTEDEAIYFCALWYSNHWVFGGGTKLTVLGQP |
| 1MFD | QAVVTQESALTTSPGETVTLTCRSSTG**TV**TSGNHANWVQEKPDHLFTGLIGDTNNRAPGVPARFSGSLIGDKAALTITGAQPEDEAIYFCALWCNNHWIFGGGTKLTVLGQP |
| **6BE2** | QLVLTQSPSASASLGASVKLTCTLSSG**HS**NYAIAWHQQQPGKGPRYLMKVNRDGSHIRGDGIPDRFSGSTSGAERYLTISSLQSEDEADYYCQTWGAGIRVFGGGTKLTVLGQP |
| **Kappa Light Chains** | |
| 2ZKH | QVVLTQSPGIMSASPGEKVTITCSASSS**VS**YMYWFQQKPGTSPKLWIYSTSNLASGVPARFRGSGSGTSYSLTISRMEAEDAATYYCQQRSGYPRTFGGGTKLEIKRA |
| 6TCM | DIQLTQSPSSLSASVGDRVTITCRASQS**VD**YDGDSYMNWYQQKPGKAPKLLIYAASYLESGVPSRFSGSGSGTDFTLTISSLQPEDFATYYCQQSHEDPYTFGQGTKVEIKRTV |
| 6Z7X | DIQMTQSPSSLSASLGGRVTITCKASQD**IN**KYLAWYQHKPGKGPRLLIHYTSTLQPGIPSRFSGSGSGRDYSFSISNLEPEDVATYYCLQYDSLLSFGAGTKLELKRA |
| 5VH3 | DILLTQSPAILSVSPGERVSFSCRASQF**VG**SSIHWYQQRTNGSPRLLIKYASESMSGIPSRFSGSGSGTDFTLSINTVESEDIADYYCQQSHSWPFTFGSGTNLEVKRTV |
| 1CR9 | DVVMTQTPLSLSVTIGQPASISCKSSQS**LL**DSDGKTYLIWVFQRPGQSPKRLIFLVSKRDSGVPDRFTGSGSGTDFTLKISRVEAEDVGVYYCWQGTHFPHTVGGGTKLEIARAD |
| 5L88 | DIVLTQTPAIMSASLGERVTMTCTANSS**VS**SNYFHWYQQKPGSSPKLWIYSTSNLASGVPTRFSGSGSGTSYSLTLSSMEAEDAATYYCHQYHRSPPTFGSGTKLKMKRA |
| 5OPY | ELVMTQTPATLSVTPGDSVSLSCRASQS**IS**NHLHWYQQKSHESPRLLIKYASQSISGIPSRFSGSGSGTDFTLSINSVETEDFGMYFCQQSNSWPHTFGGGTKLEIKRA |
| 4M43 | ELVMTQSPAILSVSPGERVSFSCRASQI**IG**TSIHWYQQRTNGSPRLLIKYASESISGIPSRFSGSGSGTDFTLTINSVESDDIADYYCQQSNSWPVTFGAGTKLELKRA |

**SUPPLIEMENTARY TABLE 6**

Table of Antibody Template Protein Light Chain Amino Acid Sequences

**Supplemental Table 6.** The PDB ID and the amino acid sequences of eight lambda light chains and eight kappa light chains are used for pivot point structural and sequential analysis. The L29 and H30 residues (IMGT numbering scheme) are in **bold** and colored in red. The L30 residues (IMGT numbering scheme) are in **bold** and colored in orange. The 6BE2 sequence is also highlighted due to the fact that it has no pivot point feature. This sequence was later found to originate from a variable domain in IgG2.

**SUPPLIEMENTARY TABLE 7a**

Table of Relative Entropy of the Residues in LCDR1 based on different numbering schemes

| Kabat | | Chothia | | AbM | | IMGT | |
| --- | --- | --- | --- | --- | --- | --- | --- |
| L23 | 3.842 | L23 | 3.842 | L23 | 3.842 | 23 | 3.889 |
| L24 | 1.084 | L24 | 1.084 | L24 | 1.084 | 24 | 1.084 |
| L25 | 1.084 | L25 | 1.084 | L25 | 1.084 | 25 | 1.084 |
| L26 | 1.341 | L26 | 1.341 | L26 | 1.341 | 26 | 1.341 |
| L27 | 1.199 | L27 | 1.199 | L27 | 1.199 | 27 | 1.199 |
| L27a | 1.264 | L28 | 0.856 | L28 | 0.856 | 28 | 0.860 |
| **L27b** | **1.126** | **L29** | **0.938** | **L29** | **0.938** | **29** | **0.984** |
| L27c | 1.075 | L30 | 0.507 | L30 | 0.507 | 30 | 0.951 |
| L27d | - | L30a | 0.752 | L30a | 0.752 | 31 | 0.994 |
| L27e | - | L30b | 0.675 | L30b | 0.675 | 32 | - |
| L27f | - | L30c | 0.711 | L30c | 0.711 | 33 | - |
| L28 | 0.593 | L30d | - | L30d | - | 34 | - |
| L29 | 0.710 | L30e | - | L30e | - | 35 | 0.969 |
| L30 | 0.580 | L30f | - | L30f | - | 36 | 0.657 |
| L31 | 0.980 | L31 | 1.023 | L31 | 1.023 | 37 | 0.993 |
| L32 | 1.158 | L32 | 1.173 | L32 | 1.173 | 38 | 1.158 |
| L33 | 1.389 | L33 | 1.443 | L33 | 1.426 | 39 | 1.389 |
| L34 | 0.940 | L34 | 0.940 | L34 | 0.940 | 40 | 0.940 |
| L35 | 4.141 | L35 | 4.141 | L35 | 4.141 | 41 | 3.356 |

**Supplemental Table 7a.** The relative entropy of conservation calculated in LCDR1 loop. Non-CDR residues, as defined by each of the numbering schemes, are highlighted in blue. The pivot point residues (L27b in Kabat and L29 in other schemes) are emphasized in bold. Residues for which insufficient sequential statistical information was available are shaded in grey (<30% occupancy). Additionally, highly conserved Cysteine and Tyrosine residues located at the termini of the LCDR1 have been color-coded in yellow and magenta, respectively.

**SUPPLIEMENTARY TABLE 7b**

Table of Relative Entropy of the Residues in HCDR1 based on different numbering schemes

| Kabat | | Chothia | | AbM | | IMGT | |
| --- | --- | --- | --- | --- | --- | --- | --- |
| H22 | 3.938 | H22 | 3.938 | H22 | 3.938 | 23 | 3.938 |
| H23 | 1.241 | H23 | 1.241 | H23 | 1.241 | 24 | 1.241 |
| H24 | 1.544 | H24 | 1.544 | H24 | 1.544 | 25 | 1.544 |
| H25 | 1.776 | H25 | 1.776 | H25 | 1.776 | 26 | 1.776 |
| H26 | 1.835 | H26 | 1.835 | H26 | 1.835 | 27 | 1.835 |
| H27 | 1.518 | H27 | 1.518 | H27 | 1.518 | 28 | 1.518 |
| H28 | 1.192 | H28 | 1.192 | H28 | 1.192 | 29 | 1.192 |
| **H29** | **2.192** | **H29** | **2.192** | **H29** | **2.192** | **30** | **2.267** |
| H30 | 0.951 | H30 | 0.951 | H30 | 0.951 | 31 | 0.896 |
| H31 | 0.839 | H31 | 0.839 | H31 | 0.839 | 32 | 0.776 |
| H32 | 1.337 | H31a | - | H31a | - | 32.1 | - |
| H33 | 0.582 | H31b | - | H31b | - | 32.2 | - |
| H34 | 1.998 | H31c | - | H31c | - | 35 | 0.896 |
| H35 | 1.271 | H31d | - | H31d | - | 36 | 0.776 |
| H32 | 1.337 | H32 | 1.597 | H32 | 1.597 | 37 | 1.597 |
| H33 | 0.582 | H33 | 0.706 | H33 | 0.706 | 38 | 0.706 |
| H34 | 1.998 | H34 | 2.298 | H34 | 2.298 | 39 | 2.298 |
| H35 | 1.271 | H35 | 1.443 | H35 | 1.443 | 40 | 1.443 |
| H36 | 3.363 | H36 | 3.363 | H36 | 3.363 | 41 | 3.363 |

**Supplemental Table 7b.** The relative entropy of conservation calculated in HCDR1 loop. Non-CDR residues, as defined by each of the numbering schemes, are highlighted in blue. The pivot point residues (H30 in IMGT and H29 in other schemes) are emphasized in **bold**. Residues for which insufficient sequential statistical information was available are shaded in grey. Additionally, highly conserved Cysteine and Tyrosine residues located at the termini of the LCDR1 have been color-coded in yellow and magenta, respectively. Note that in Kabat numbering scheme, the insertion site was placed at H35 as no residue was equivalent to the H31a-d residues. Additionally, in the IMGT numbering scheme, the H33 and H34 residues lack equivalent residues in the other numbering schemes. These residues also lack sufficient database information, with both residues having <30% occupancy.

**SUPPLIEMENTARY TABLE 8a**

Table of the Average Relative Entopy of Differences Calculated in each Regions in the Variable Domains

| **Region** | **Avg. RE** (Differences) | **Region** | **Ave. RE** (Differences) |
| --- | --- | --- | --- |
| LFR-1 | 0.496 | HFR-1 | 0.396 |
| LFR-2 | 0.395 | HFR-2 | 0.425 |
| LFR-3 | 0.396 | HFR-3 | 0.517 |
| LFR-4 | 0.456 | HFR-4 | 0.219 |
| LCDR1 | 0.754 | HCDR1 | 0.437 |
| LCDR2 | 0.676 | HCDR2 | 0.561 |
| LCDR3 | 0.884 | HCDR3 | 0.315 |
| L-Frameworks | 0.432 | H-Frameworks | 0.422 |
| LCDRs | 0.776 | HCDRs | 0.450 |
| Light Chain | 0.520 | Heavy Chain | 0.431 |

**Supplemental Table 8a.** The average relative entropy of differences (Avg.RE (Differences)) calculated in different regions (Kabat Numbering scheme) for the Homo Sapiens immunoglobulin variable domains.

**SUPPLIEMENTARY TABLE 8b**

Table of the Average, Median, Standard Deviation of the Relative Entopy of Differences Calculated and outliers in each Regions in the Variable Domains

| **Region** | **Avg. RE**  (Differences) | **Mdn. RE**  (Differences) | **Std. RE**  (Differences) | **Outliers** (RE.D) |
| --- | --- | --- | --- | --- |
| LFR-1 | 0.496 | 0.324 | 0.526 | L2 (2.026) |
|  |  |  |  | L11 (1.397) |
| LFR-2 | 0.395 | 0.166 | 0.654 | L43 (2.658) |
| LFR-3 | 0.396 | 0.165 | 0.506 | L66 (2.261) |
|  |  |  |  | L81 (1.341) |
| LFR-4 | 0.456 | 0.193 | 0.595 | L100 (1.832) |
| HFR-1 | 0.396 | 0.223 | 0.369 | H5 (1.104) |
|  |  |  |  | H19 (1.184) |
| HFR-2 | 0.425 | 0.176 | 0.561 | H40 (1.853) |
| HFR-3 | 0.506 | 0.430 | 0.517 | H67 (1.815) |
|  |  |  |  | H75 (1.632) |
|  |  |  |  | H84 (1.545) |
| HFR-4 | 0.219 | 0.074 | 0.304 | H108 (0.952) |
| LCDR1 | 0.754 | 0.438 | 0.935 | L27B (3.394) |
| LCDR2 | 0.676 | 0.327 | 0.663 | L51 (1.511) |
|  |  |  |  | L55 (1.757) |
| LCDR3 | 0.884 | 0.768 | 0.553 | L90 (2.066) |
| HCDR1 | 0.437 | 0.259 | 0.403 | H27 (1.348) |
| HCDR2 | 0.561 | 0.515 | 0.345 | H52A (1.423) |
|  |  |  |  | H60 (1.116) |
| HCDR3 | 0.315 | 0.264 | 0.278 | H102 (1.166) |

**Supplemental Table 8b.** The average (avg), median (mdn) and standard deviation (std) of the relative entropies of differences (RE.D) between mouse and human calculated in different regions (Kabat Numbering scheme) in the Homo Sapiens immunoglobulin variable domains. The residues with RE.D value higher than the average RE.D plus 1.65 times the standard deviation were identified as the outliers in the group.

**SUPPLIEMENTARY TABLE 9**

Table of the Amino Acid Distributions at the Outlier Residues in Human and Mouse

| **Region** | **Residue** | **RE**  **(Diff.)** | **Homo sapiens AA Distribution** | **Mus Musculus AA Distribution** |
| --- | --- | --- | --- | --- |
| LFR-1 | L2 | 2.026 | I (42%), S (34%), Y (8%) | I (71%), V (15%) A (6%) |
|  | L11 | 1.397 | L (53%), V (37%), A (7%) | L (63%), M (31%) |
| LCDR1 | L27B | 3.394 | N (45%), D(19%),  L (14%), V (9%), | L (50%), V (20%).  I (14%), A (13%) |
| LFR-2 | L43 | 2.658 | A (76%), S (10%), P (8%) | S (65%), P (15%), T (9%) |
| LCDR2 | L51 | 1.511 | A (42%), N (20%),  D (11%), V (11%), T (6%) | T (39%), A (37%), V (16%) |
|  | L55 | 1.757 | P (42%), A (23%),  Q (15%), E (10%), | A (35%), E (18%), F (13%),  H (7%), Y (6%), D (6%) |
| LFR-3 | L66 | 2.261 | G (51%), K (30%) | G (89%), L (6%) |
|  | L83 | 1.341 | E (45%), F (37%), V (12%) | L (33%), A (26%),  V (9%), I (8%), E (6%) |
| LCDR3 | L90 | 2.066 | Q (47%), S (17%),  T (14%), A (8%), | Q (79%), L (7%),  H (6%), N (6%), |
| LFR-4 | L100 | 1.832 | G (36%), Q (33%), T (17%) | G (62%), A (22%), S (13%) |
| HFR-1 | H5 | 1.104 | V (52%), L (19%),  Q (17%), E (5%), | Q (79%),V (10%) |
|  | H19 | 1.184 | R (52%), K (22%), S (20%) | K (88%), S (13%) |
| HCDR1 | H27 | 1.348 | F (48%), Y (18%),  G (17%), D (7%), | Y (66%), F (30%) |
| HFR-2 | H40 | 1.853 | A (65%), P (17%), S (6%) | R (56%), S (13%),  T (9%), P (9%), A (6%) |
| HCDR2 | H52A | 1.423 | P (27%), G (17%)  Y (15%), S (13%), A (6%), | P (79%), S (9%) |
|  | H60 | 1.116 | A (60%), N (16%), S (13%) | N (72%) A (9%), P (6%) |
| HFR-3 | H67 | 1.815 | F (47%), V (40%),  L (6%), I (5%), | A (66%), F (18%), L (8%) |
|  | H75 | 1.632 | K (62%), T (12%), I (9%) | S (68%), K (23%) |
|  | H84 | 1.545 | A (53%), S (19%),  P (10%), V (5%) | S (77%), T (12%), A (6%) |
| HCDR3 | H102 | 1.166 | Y (33%), V (24%),  I (9%), P (8%), L (8%) | Y (79%), V (14%) |
| HFR-4 | H108 | 0.952 | L (63%), T (22%), M (9%) | T (53%), S (24%), L (21%) |

**Supplemental Table 9.** The amino acid frequencies ( > 5%) of the outliers based on the relative entropy differences between the mouse and the human immunoglobulin variable domain.

**SUPPLIEMENTARY TABLE 10**

Table of the Residue Relative Entropy, Consensus Mouse variable domain Sequences and Calculated AbNatiV Score based on the consensus sequence

| **VL** | **H_C_** | **M_C_** | **RE** | **(1-AbN Score) kappa** | **(1-AbN Score) lambda** | **VH** | **H_C_** | **M_C_** | **RE** | **(1-AbN Score)** |
| --- | --- | --- | --- | --- | --- | --- | --- | --- | --- | --- |
| L1 | **D** | **D** | 0.518 | 0.000 | **0.076** | H1 | **Q** | **Q** | 0.070 | 0.000 |
| L2 | **I** | **I** | **2.026** | 0.000 | **0.076** | H2 | **V** | **V** | 0.120 | 0.000 |
| L3 | **V** | **V** | 0.471 | **0.072** | 0.000 | H3 | **Q** | **Q** | 0.185 | 0.000 |
| L4 | L | M | 0.195 | 0.000 | **0.088** | H4 | **L** | **L** | 0.018 | 0.000 |
| L5 | **T** | **T** | 0.055 | 0.000 | 0.000 | H5 | V | Q | **1.104** | **0.091** |
| L6 | **Q** | **Q** | 0.045 | 0.000 | 0.000 | H6 | E | Q | 0.223 | 0.000 |
| L7 | **S** | **S** | 1.221 | 0.000 | **0.040** | H7 | **S** | **S** | 0.659 | 0.000 |
| L8 | **P** | **P** | 0.167 | 0.000 | **0.091** | H8 | **G** | **G** | 0.012 | 0.000 |
| L9 | S | A | 0.934 | **0.084** | **0.073** | H9 | G | A | 0.452 | 0.000 |
| L10 | **S** | **S** | 0.593 | 0.000 | 0.001 | H10 | G | E | 0.496 | 0.000 |
| L11 | **L** | **L** | **1.397** | 0.000 | 0.000 | H11 | **L** | **L** | 0.998 | **0.091** |
| L12 | **S** | **S** | 0.322 | 0.000 | 0.000 | H12 | **V** | **V** | 0.689 | **0.089** |
| L13 | A | V | 1.254 | 0.003 | 0.000 | H13 | **K** | **K** | 0.369 | 0.000 |
| L14 | **S** | **S** | 0.430 | **0.041** | **0.032** | H14 | **P** | **P** | 0.008 | 0.000 |
| L15 | P | L | 0.383 | **0.064** | **0.069** | H15 | **G** | **G** | 0.094 | 0.000 |
| L16 | **G** | **G** | 0.010 | 0.000 | 0.000 | H16 | G | A | 0.915 | 0.000 |
| L17 | Q | E | 0.299 | **0.058** | **0.055** | H17 | **S** | **S** | 0.188 | 0.001 |
| L18 | **R** | **R** | 0.332 | 0.000 | **0.064** | H18 | L | V | 0.598 | 0.000 |
| L19 | **V** | **V** | 0.074 | 0.000 | 0.005 | H19 | **K** | **K** | **1.184** | 0.000 |
| L20 | **T** | **T** | 0.293 | 0.000 | 0.000 | H20 | **L** | **L** | 0.462 | **0.021** |
| L21 | **I** | **I** | 0.324 | 0.000 | 0.000 | H21 | **S** | **S** | 0.044 | 0.000 |
| L22 | S | T | 0.065 | 0.000 | 0.000 | H22 | **C** | **C** | 0.002 | 0.000 |
| L23 | **C** | **C** | 0.007 | 0.000 | 0.000 | H23 | A | K | 0.683 | 0.003 |
| L24 | **R** | **R** | 0.287 | 0.000 | **0.059** | H24 | **A** | **A** | 0.139 | 0.000 |
| L25 | **A** | **A** | 1.941 | 0.000 | **0.082** | H25 | **S** | **S** | 0.181 | 0.000 |
| L26 | **S** | **S** | 0.455 | 0.000 | **0.019** | H26 | **G** | **G** | 0.108 | 0.000 |
| L27 | **Q** | **Q** | 0.288 | 0.002 | **0.089** | H27 | F | Y | **1.348** | 0.000 |
| L27A | **S** | **S** | 0.053 | **0.015** | 0.001 | H28 | **T** | **T** | 0.133 | 0.000 |
| L27B | N | L | **3.394** | 0.000 | 0.001 | H29 | **F** | **F** | 0.262 | 0.000 |
| L28 | S | N | 1.138 | 0.001 | 0.055 | H30 | S | T | 0.860 | 0.001 |
| L29 | **G** | **G** | 0.438 | **0.088** | **0.046** | H31 | **S** | **S** | 0.125 | 0.000 |
| L30 | **S** | **S** | 0.270 | **0.090** | **0.090** | H32 | **Y** | **Y** | 0.180 | 0.000 |
| L31 | N | S | 0.390 | 0.000 | 0.000 | H33 | **W** | **W** | 0.592 | **0.058** |
| L32 | **Y** | **Y** | 0.159 | 0.000 | 0.000 | H34 | **M** | **M** | 0.255 | 0.000 |
| L33 | **L** | **L** | 0.490 | 0.000 | **0.052** | H35 | **H** | **H** | 0.506 | 0.000 |
| L34 | A | H | 0.507 | 0.002 | **0.043** | H36 | **W** | **W** | 0.002 | 0.000 |
| L35 | **W** | **W** | 0.002 | 0.000 | 0.000 | H37 | V | **V** | 0.157 | 0.000 |
| L36 | **Y** | **Y** | 0.124 | 0.001 | **0.018** | H38 | R | K | 1.130 | **0.088** |
| L37 | **Q** | **Q** | 0.123 | 0.000 | 0.003 | H39 | **Q** | **Q** | 0.028 | 0.000 |
| L38 | **Q** | **Q** | 0.130 | 0.000 | 0.000 | H40 | A | R | **1.853** | **0.085** |
| L39 | **K** | **K** | 0.351 | 0.000 | 0.000 | H41 | **P** | **P** | 0.128 | 0.000 |
| L40 | **P** | **P** | 0.159 | 0.000 | 0.000 | H42 | **G** | **G** | 0.313 | 0.000 |
| L41 | **G** | **G** | 0.198 | 0.000 | 0.000 | H43 | **K** | **K** | 0.314 | 0.000 |
| L42 | **Q** | **Q** | 0.360 | 0.000 | 0.000 | H44 | **G** | **G** | 0.194 | 0.005 |
| L43 | A | S | **2.658** | **0.012** | **0.043** | H45 | **L** | **L** | 0.018 | 0.000 |
| L44 | **P** | **P** | 0.166 | 0.000 | 0.000 | H46 | **E** | **E** | 0.057 | 0.000 |
| L45 | **K** | **K** | 0.570 | 0.000 | **0.086** | H47 | **W** | **W** | 0.013 | 0.000 |
| L46 | **L** | **L** | 0.217 | 0.000 | 0.000 | H48 | **I** | **I** | 0.637 | **0.089** |
| L47 | **L** | **L** | 0.700 | 0.000 | 0.000 | H49 | **G** | **G** | 1.110 | 0.000 |
| L48 | **I** | **I** | 0.047 | 0.000 | 0.000 | H50 | **R** | **R** | 0.575 | **0.035** |
| L49 | **Y** | **Y** | 0.121 | 0.000 | 0.000 | H51 | **I** | **I** | 0.196 | 0.000 |
| L50 | G | Y | 0.436 | **0.054** | **0.066** | H52 | S | D | 0.705 | 0.000 |
| L51 | A | T | **1.511** | 0.000 | 0.004 | H52A | **P** | **P** | **1.423** | 0.000 |
| L52 | **S** | **S** | 0.194 | 0.000 | 0.000 | H53 | S | N | 0.623 | **0.063** |
| L53 | **N** | **N** | 0.327 | 0.000 | 0.002 | H54 | **G** | **S** | 0.515 | 0.000 |
| L54 | R | L | 0.294 | 0.007 | **0.088** | H55 | **G** | **G** | 0.381 | 0.000 |
| L55 | P | A | **1.757** | **0.084** | **0.089** | H56 | S | G | 0.330 | 0.006 |
| L56 | **S** | **S** | 0.214 | 0.000 | 0.000 | H57 | **T** | **T** | 0.454 | 0.000 |
| L57 | **G** | **G** | 0.043 | 0.000 | 0.000 | H58 | Y | K | 0.401 | 0.006 |
| L58 | **V** | **V** | 0.422 | 0.000 | 0.001 | H59 | **Y** | **Y** | 0.075 | 0.000 |
| L59 | **P** | **P** | 0.086 | 0.000 | 0.000 | H60 | A | N | **1.116** | **0.090** |
| L60 | **D** | **D** | 0.559 | **0.086** | 0.000 | H61 | D | E | 0.779 | 0.000 |
| L61 | **R** | **R** | 0.019 | 0.000 | 0.000 | H62 | S | K | 0.894 | 0.000 |
| L62 | **F** | **F** | 0.010 | 0.000 | 0.000 | H63 | V | F | 0.526 | 0.000 |
| L63 | **S** | **S** | 0.173 | 0.000 | 0.000 | H64 | **K** | **K** | 0.309 | **0.090** |
| L64 | **G** | **G** | 0.046 | 0.000 | 0.000 | H65 | **G** | **G** | 0.235 | 0.000 |
| L65 | **S** | **S** | 0.016 | 0.000 | 0.000 | H66 | R | K | 1.142 | **0.091** |
| L66 | **G** | **G** | **2.261** | 0.000 | **0.042** | H67 | F | A | **1.815** | **0.073** |
| L67 | **S** | **S** | 0.114 | 0.000 | 0.001 | H68 | **T** | **T** | 0.041 | 0.000 |
| L68 | **G** | **G** | 0.072 | 0.000 | 0.000 | H69 | I | L | 0.858 | 0.000 |
| L69 | **T** | **T** | 0.421 | 0.000 | 0.000 | H70 | S | T | 0.597 | 0.000 |
| L70 | **D** | **D** | 1.113 | 0.000 | **0.090** | H71 | R | V | 0.609 | **0.081** |
| L71 | **F** | **F** | 0.812 | 0.000 | **0.058** | H72 | **D** | **D** | 0.021 | 0.001 |
| L72 | **T** | **T** | 0.148 | 0.000 | 0.002 | H73 | N | K | 0.538 | **0.055** |
| L73 | **L** | **L** | 0.008 | 0.000 | 0.000 | H74 | **S** | **S** | 0.292 | 0.000 |
| L74 | **T** | **T** | 1.079 | 0.000 | 0.000 | H75 | K | S | **1.632** | **0.077** |
| L75 | **I** | **I** | 0.029 | 0.000 | 0.000 | H76 | N | S | 0.626 | 0.000 |
| L76 | **S** | **S** | 0.175 | 0.000 | 0.000 | H77 | **T** | **T** | 0.215 | 0.000 |
| L77 | G | S | 0.636 | 0.000 | **0.022** | H78 | **A** | **A** | 0.556 | 0.003 |
| L78 | L | V | 0.946 | **0.089** | 0.009 | H79 | **Y** | **Y** | 0.827 | 0.000 |
| L79 | Q | E | 0.448 | 0.000 | **0.063** | H80 | L | M | 0.412 | 0.000 |
| L80 | **A** | **A** | 0.453 | 0.000 | 0.000 | H81 | **Q** | **Q** | 0.310 | **0.090** |
| L81 | **E** | **E** | 0.521 | 0.000 | 0.000 | H82 | **L** | **L** | 0.448 | 0.000 |
| L82 | **D** | **D** | 0.005 | 0.000 | 0.000 | H82A | N | S | 0.272 | 0.000 |
| L83 | E | L | **1.341** | **0.030** | **0.091** | H82B | **S** | **S** | 0.098 | 0.000 |
| L84 | **A** | **A** | 0.112 | 0.081 | 0.000 | H82C | **L** | **L** | 0.206 | 0.000 |
| L85 | D | T | 0.451 | 0.000 | **0.082** | H83 | **T** | **T** | 0.773 | 0.000 |
| L86 | **Y** | **Y** | 0.001 | 0.000 | 0.000 | H84 | A | S | **1.545** | 0.000 |
| L87 | **Y** | **Y** | 0.157 | 0.000 | 0.000 | H85 | **E** | **E** | 0.610 | 0.000 |
| L88 | **C** | **C** | 0.001 | 0.000 | 0.000 | H86 | **D** | **D** | 0.006 | 0.000 |
| L89 | **Q** | **Q** | 0.423 | 0.008 | **0.025** | H87 | T | S | 0.970 | **0.090** |
| L90 | **Q** | **Q** | **2.066** | 0.002 | **0.055** | H88 | **A** | **A** | 0.003 | 0.000 |
| L91 | **Y** | **Y** | 0.342 | 0.001 | **0.061** | H89 | **V** | **V** | 0.063 | 0.000 |
| L92 | D | S | 0.754 | 0.000 | **0.085** | H90 | **Y** | **Y** | 0.003 | 0.000 |
| L93 | **S** | **S** | 0.382 | 0.000 | 0.000 | H91 | **Y** | **Y** | 0.011 | 0.000 |
| L94 | S | Y | 0.863 | 0.002 | 0.001 | H92 | **C** | **C** | 0.000 | 0.000 |
| L95 | **P** | **P** | 1.387 | 0.000 | 0.000 | H93 | **A** | **A** | 0.024 | 0.000 |
| L96 | Y | L | 0.972 | 0.000 | **0.010** | H94 | **R** | **R** | 0.276 | 0.000 |
| L97 | **T** | **T** | 0.768 | 0.000 | **0.090** | H95 | D | Y | 0.394 | **0.050** |
| L98 | **F** | **F** | 0.019 | 0.000 | 0.000 | H96 | G | Y | 0.264 | 0.003 |
| L99 | **G** | **G** | 0.003 | 0.000 | 0.000 | H97 | **Y** | **Y** | 0.286 | 0.002 |
| L100 | **G** | **G** | **1.832** | 0.000 | 0.000 | H98 | G | Y | 0.257 | **0.033** |
| L101 | **G** | **G** | 0.007 | 0.000 | 0.000 | H99 | G | Y | 0.124 | 0.000 |
| L102 | **T** | **T** | 0.003 | 0.000 | 0.000 | H100 | G | Y | 0.122 | **0.060** |
| L103 | **K** | **K** | 0.146 | 0.000 | 0.000 | H100A | **Y** | **Y** | 0.223 | 0.000 |
| L104 | **L** | **L** | 1.218 | 0.001 | 0.000 | H100B | Y | F | 0.330 | **0.075** |
| L105 | **E** | **E** | 0.832 | 0.000 | **0.090** | H100C | Y | Y | 0.410 | **0.030** |
| L106 | **I** | **I** | 0.793 | 0.000 | **0.091** | H101 | **D** | **D** | 0.216 | 0.000 |
| L107 | **K** | **K** | 0.239 | 0.000 | **0.088** | H102 | **Y** | **Y** | **1.166** | 0.000 |
| L108 | **R** | **R** | 0.376 | **0.087** | **0.087** | H103 | **W** | **W** | 0.003 | 0.000 |
|  |  |  |  |  |  | H104 | **G** | **G** | 0.002 | 0.000 |
|  |  |  |  |  |  | H105 | **Q** | **Q** | 0.419 | 0.000 |
|  |  |  |  |  |  | H106 | **G** | **G** | 0.004 | 0.000 |
|  |  |  |  |  |  | H107 | **T** | **T** | 0.074 | 0.000 |
|  |  |  |  |  |  | H108 | L | T | **0.952** | **0.033** |
|  |  |  |  |  |  | H109 | **V** | **V** | 0.536 | 0.000 |
|  |  |  |  |  |  | H110 | **T** | **T** | 0.162 | 0.000 |
|  |  |  |  |  |  | H111 | **V** | **V** | 0.016 | 0.000 |
|  |  |  |  |  |  | H112 | **S** | **S** | 0.017 | 0.000 |
|  |  |  |  |  |  | H113 | **S** | **S** | 0.227 | 0.000 |

**Supplemental Table 10.** The Relative Entropy calculated based on the amino acid distribution differences between Homo Sapien and Mus antibody variable domain sequences. The amino acid consensus Human (Hc) and Mouse (Mc) were listed, with identical sequences in **bold**. The AbNatiV Score (AbN score) of the consensus Mus amino acid sequences represent the humanness score of each residue based on VL kappa, VL lambda and VH models. To align with the relative entropy, we used (1-AbNatiV Score) to depict the nonhumanness level of the sequences. The hotspots identified in the RE analysis and the AbNatiV analysis ((1-AbNative Score) > 0.01) were highlighted in **red**. The CDRs (Kabat definition) were highlighted in red, green and magenta respectively.

**SUPPLIEMENTARY TABLE 11**

|  | **Yeast** | **Variable Domains** | **Variable Light**  **(VL)** | **Variable Heavy**  **(VH)** | **Homo**  **Sapiens**  **Variable Domain** | **Mus**  **Musculus**  **Variable Domain** |
| --- | --- | --- | --- | --- | --- | --- |
| A | 5.62 | 6.45 | 6.11 | 6.58 | 6.50 | 6.18 |
| C | 1.29 | 1.91 | 1.98 | 1.89 | 1.90 | 1.84 |
| D | 5.78 | 4.32 | 4.14 | 4.39 | 4.49 | 4.05 |
| E | 6.48 | 3.28 | 3.04 | 3.37 | 3.07 | 3.72 |
| F | 4.45 | 3.09 | 3.26 | 3.03 | 3.14 | 3.16 |
| G | 5.06 | 10.26 | 10.08 | 10.34 | 10.49 | 9.33 |
| H | 2.14 | 0.81 | 0.81 | 0.80 | 0.78 | 0.98 |
| I | 6.51 | 3.53 | 4.51 | 3.15 | 3.58 | 3.23 |
| K | 7.27 | 4.29 | 3.70 | 4.52 | 3.74 | 5.88 |
| L | 9.50 | 7.05 | 7.22 | 6.99 | 7.04 | 7.02 |
| M | 2.09 | 1.58 | 0.76 | 1.90 | 1.56 | 1.91 |
| N | 6.05 | 2.97 | 2.63 | 3.10 | 2.97 | 2.94 |
| P | 4.39 | 3.83 | 5.71 | 3.11 | 3.83 | 3.77 |
| Q | 3.94 | 5.03 | 6.36 | 4.52 | 5.07 | 4.94 |
| R | 4.42 | 4.43 | 3.83 | 4.66 | 4.97 | 3.53 |
| S | 8.90 | 13.14 | 14.72 | 12.53 | 13.09 | 13.02 |
| T | 5.88 | 8.24 | 8.78 | 8.03 | 7.83 | 8.65 |
| V | 5.65 | 6.56 | 5.50 | 6.98 | 6.87 | 5.67 |
| W | 1.04 | 2.84 | 1.52 | 3.35 | 2.92 | 2.97 |
| Y | 3.36 | 6.34 | 5.34 | 6.72 | 6.17 | 7.20 |

Table of Antibody Variable Domains Amino Acids Distribution and the Codon Usage in Yeast

**Supplemental Table 11.** Amino acid frequency calculated based on codon usage (yeast) and sequence stored in the abYsis database (Variable Domains, Variable Light Domain, Variable Heavy Domain, Homo Sapiens Variable Domain, Mus Musculus Variable Domain).
